# Analysis of non-coding RNAs in *Methylorubrum extorquens* reveals a novel small RNA specific to Methylobacteriaceae

**DOI:** 10.1101/2022.01.24.477521

**Authors:** Emilie Boutet, Samia Djerroud, Katia Smail, Marie-Josée Lorain, Meiqun Wu, Martin Lamarche, Roqaya Imane, Carlos Miguez, Jonathan Perreault

## Abstract

*Methylorubrum extorquens* metabolizes methanol, a cheap raw material that can be derived from waste. It is a facultative methylotroph, making it a model organism to study the metabolism of one carbon compounds. Despite a considerable interest to exploit this bacteria as a biotechnological tool in a methanol-based bioeconomy, little is known about its non-coding sRNA. Small RNAs play well-documented essential roles in *Escherichia coli* for post-transcriptional regulation; and have important functions in many bacteria, including other Alphaproteobacteria like *Agrobacterium tumefaciens. M. extorquens* is expected to contain many sRNAs, especially since it also encodes for the protein Hfq, a chaperone protein important in the interaction between sRNAs and their target, but also critical for the stabilization of sRNAs themselves. Few sRNAs are annotated in the genome of this Alphaproteobacteria and they were never validated. In this study, formerly annotated sRNAs ffh, CC2171, BjrC1505 were confirmed by Northern blot, validating the expression of sRNAs in *M. extorquens*. Moreover, analysis of RNA-sequencing data established a considerable list of potential sRNAs. Interesting candidates selected after bioinformatic analysis were tested by Northern blot, revealing a novel sRNA specific to Methylobacteriaceae, sRNA Met2624. Its expression patterns and genomic context were analyzed. This research is the first experimental validation of sRNAs in *M. extorquens* and paves the way for other sRNA discoveries.

---

*Methylorubrum extorquens* (formerly *Methylobacterium extorquens*) [^1^] has potential in a future C1-carbon based bioeconomy for its ability to produce value-added product at a large scale from a cheap raw material derived from waste, methanol (and other C1 compounds, i.e. with a single carbon). This Alphaproteobacteria has already been engineered to produce numerous value-added products from methanol including heterologous proteins like insecticidal protein (Cry1Aa) [^2, 3^], metabolites from the ethylmalonyl-CoA (EMC) pathway [^4-8^] and metabolites from the polyhydroxybutyrate (PHB) cycle [^9-13^]. *M. extorquens* is also a model organism for the study of C1 consumption, since it can also use other carbon sources to grow [^14^]. Knockout mutants can therefore be produced in genes essential for C1 consumption without impeding the ability of the bacteria to grow, making it a perfect model organism to study methylotrophy.

Despite its importance at the bioindustrial level and in fundamental research, little is known about its non-coding small RNAs (sRNA). Even though there is already profound knowledge about its metabolism, the presence of sRNAs in *M. extorquens* has not yet been studied. A better understanding of the role of sRNAs in the genetic regulation of *M. extorquens* could give hints on how to maximize industrial processes. Usually between 50 and 300 nucleotides, sRNAs are either cis or trans-encoded [^15^]. Small RNAs bind to their targets due to the complementarity of their nucleotides. This pairing affects the translation of the regulated mRNA, which can be upregulated or downregulated, more often the latter. The sRNA can bind to the ribosome binding site (RBS) for example, therefore preventing the ribosome to initiate translation. Inversely, in some cases the binding of the sRNA to its target could release a RBS that would normally be stuck within a secondary structure, therefore allowing translation to initiate [^16^]. The concurrent binding of sRNAs and the chaperone protein Hfq to their targets can also interfere with the stability of the mRNA by recruiting the ribonuclease E (RNAse E), leading to the degradation of the mRNA target ^[16-18^]. Inversely, the binding of the sRNA-Hfq complex to their target site can protect it from RNAse E cleavage, since both the chaperone protein and the nuclease interact with A/U-rich sequence contiguous to stem-loop structures [^19^].

The chaperone protein Hfq is often associated with sRNAs in Gram-negative bacteria, because it can not only stabilize the interaction between the sRNA and its target, but it can also stabilize the sRNA itself [^20, 21^]. Modes of action of sRNAs are diverse and they allow a tight regulation of genetic information. To develop *M. extorquens* has an even more powerful biotechnological tool, it is important to identify sRNAs, an efficient set of regulatory elements for gene control that this bacterium presumably already has, and perhaps that we could use to our advantage in the future. The Hfq protein is encoded within the genome of *M. extorquens* (WP_003600267.1). Jointly with this chaperone protein, sRNAs play an important role in genetic regulation in other Alphaproteobacteria of the same order (Hyphomicrobiales) such as *Brucella melitensis*, where 24 distinct sRNAs are annotated (with a E-value lower than 0.0005). *Escherichia coli*, another Proteobacteria, has approximately 80 sRNAs that were validated experimentally [^22^] out of the 103 annotated in its genome (with a E-value lower than 0.0005). Since *M. extorquens* is also a Proteobacteria that encodes for the Hfq protein, we hypothesized that it may have a similar number of sRNAs. Some sRNAs are predicted to be present in *M. extorquens*, but they were never experimentally confirmed before.

Novel noncoding RNA (ncRNA) in bacteria can be characterized by processing of RNA-sequencing (RNA-seq) data to identify highly transcribed conserved regions that are not annotated as encoding proteins. Multiple bioinformatics tools have been developed to determine interesting noncoding regions from RNA-seq data such as sRNA-Detect [^23^]. Small RNAs are discerned based on the assumptions that for a given length, reads show small coverage variation with a minimal depth coverage. We experimentally validated several sRNAs previously annotated in *M. extorquens*. Moreover, analysis of the RNA-seq data revealed numerous putative sRNAs in *M. extorquens*. We also confirmed one of them and evaluated it experimentally, describing at the same time a novel sRNA specific to Methylobacteriaceae.

## Results and discussion

### Annotated sRNAs and intergenic regions size distribution

To get a sense of the likelihood to discover sRNAs in *M. extorquens*, we first looked at the number of annotated sRNAs with an E-value lower than 0.0005 within the genomes of Alphaproteobacteria (Figure 1). All sRNAs mentioned in this study had an E-value lower than 0.0005. This information was extracted from the database RiboGap (version 2) [^24^], since it facilitates the examination of intergenic regions (IGR) from prokaryotes. Evidence about sRNAs in RiboGap comes from the Rfam database [^25^] and are limited to available covariance models used for homology searches and annotations.

**Figure 1.**
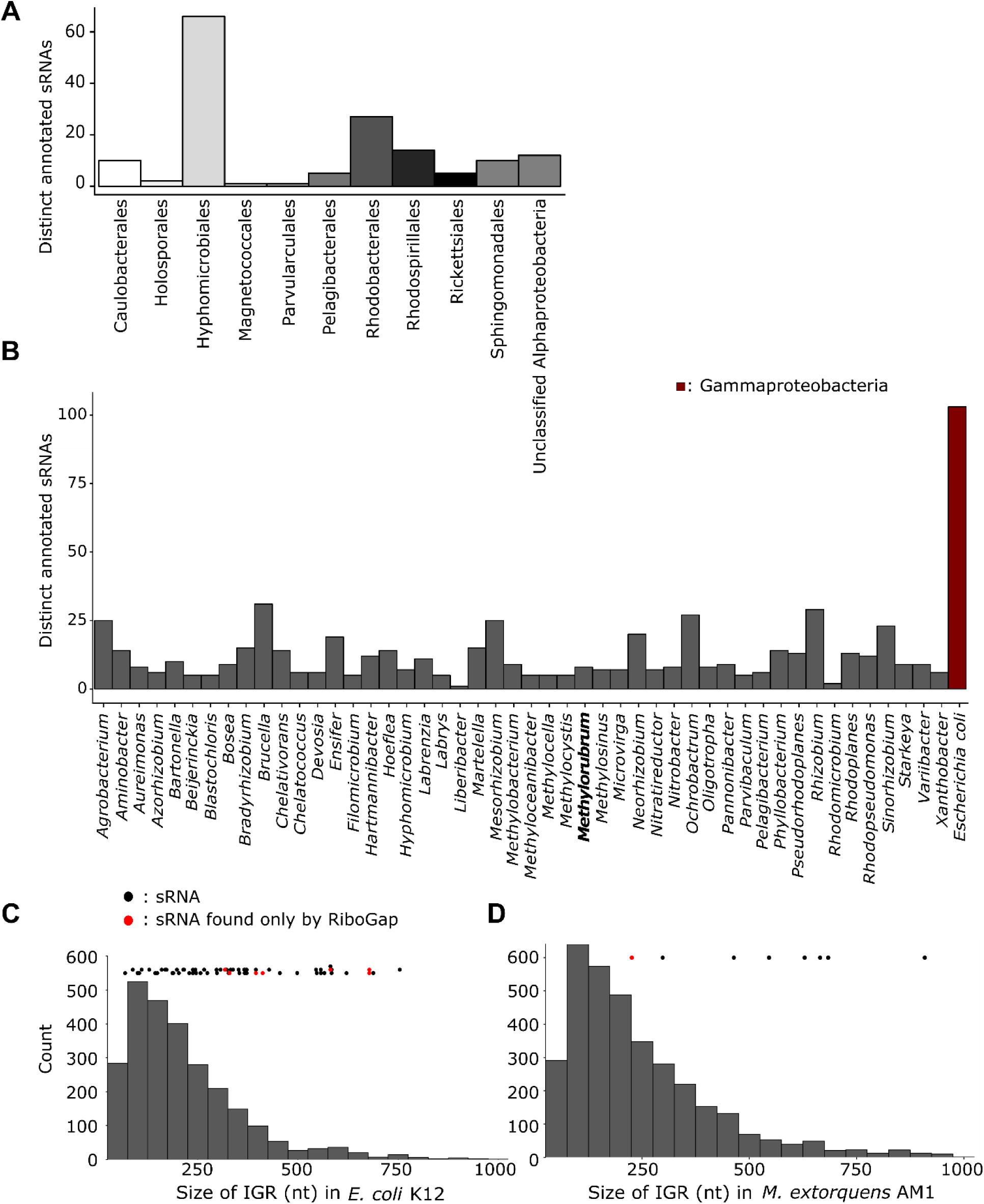
Annotated sRNAs in Alphaproteobacteria. Distinct sRNAs with an E-value smaller than 0.0005 annotated in the genome of the different orders within the Alphaproteobacteria class (A) and within different genera of the Hyphomicrobiales order (B). The different sRNAs found in the genome of the model Gammaproteobacteria *Escherichia coli* are depicted in red (A). The size distribution of intergenic gaps that are in between 50 and 1000 nucleotides for the strain *E. coli* K-12 substr. MG1655 and *M. extorquens* AM1 are shown in (C) and (D) respectively. IGRs containing a sRNA have been identified with dots, where red ones represent sRNAs identified only by RiboGap. Black dots illustrate sRNAs found by both Rfam and RiboGap database. The histograms are grouped in bins of 50.

A total of 87 distinct sRNAs were annotated in Alphaproteobacteria and spread throughout all orders of this bacterial class (Figure 1, A). The order with the highest number of distinct annotated sRNAs (66 sRNAs) is Hyphomicrobiales, which comprises *M. extorquens* (Figure 1, A). Numerous sRNAs are predicted in different genera of the order Hyphomicrobiales (Figure 1, B) where eight distinct potential sRNAs [^26-33^] are annotated in the genome of all available strains of the genus *Methylorubrum*, but they were never confirmed in laboratory conditions (Table 1). This is still very few compared to the 103 distinct sRNAs annotated within the genome of *Escherichia coli* alone (Figure 1, B), a Gammaproteobacteria model organism. All Hyphomicrobiales bacteria represented in (Figure 1, B) also encode for the chaperone protein Hfq.

**Table 1.**
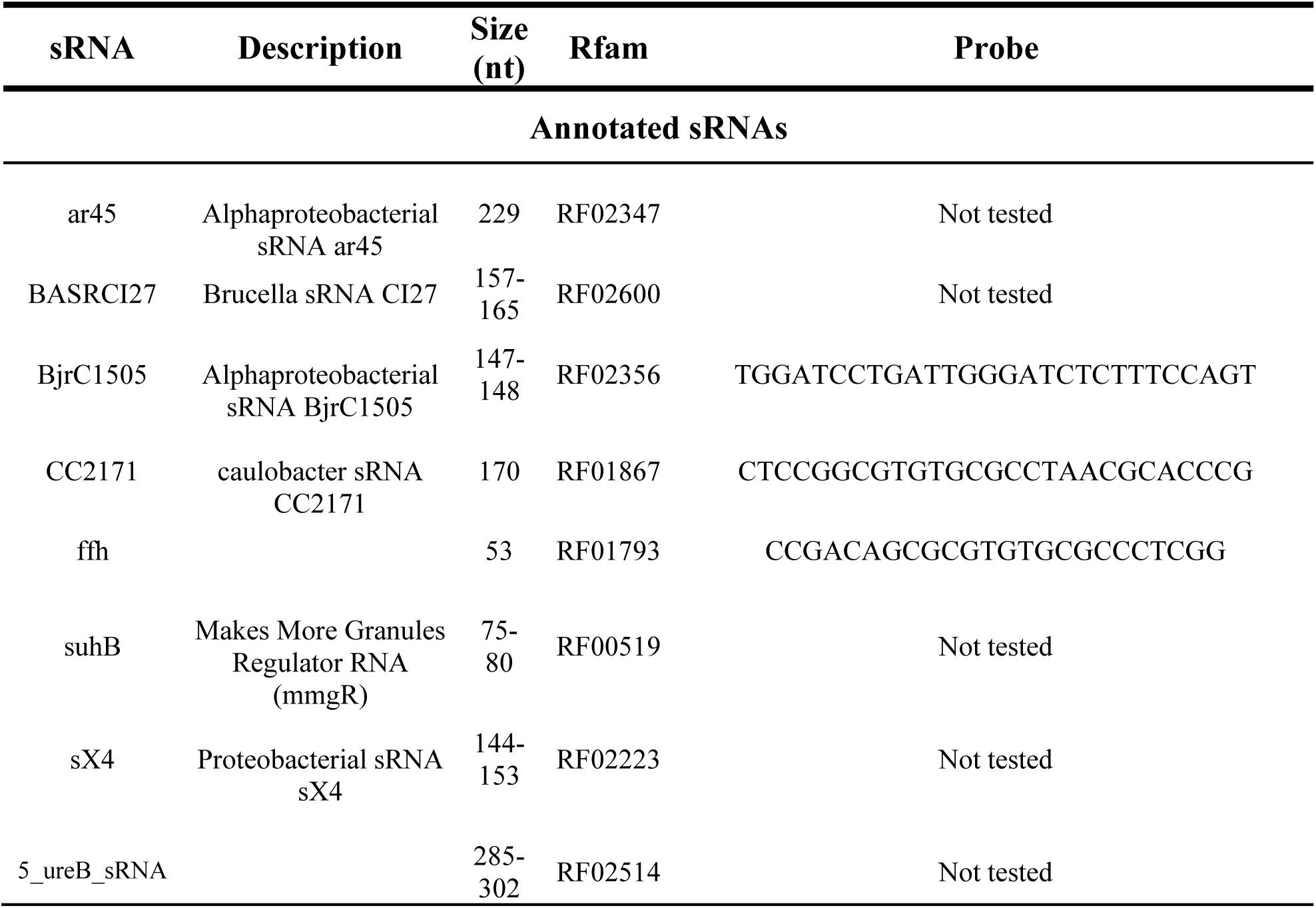
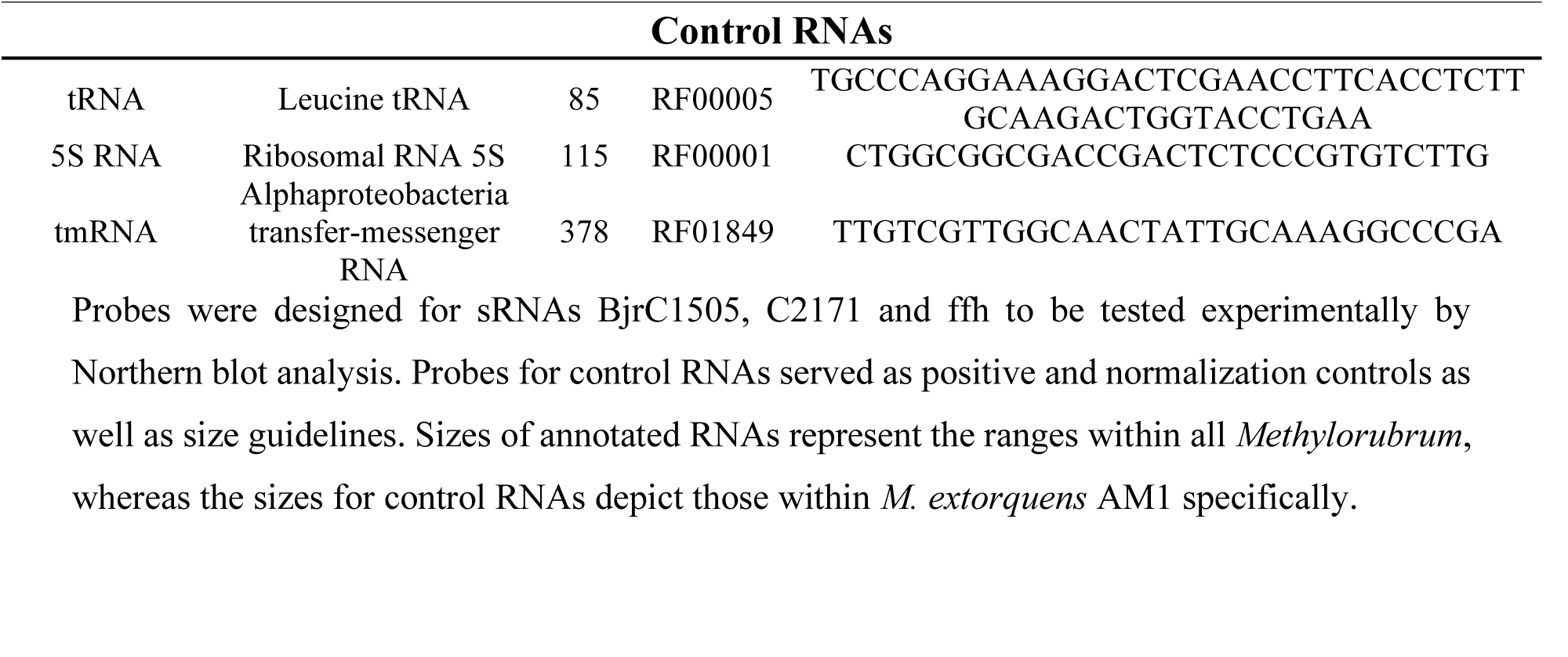
Annotated sRNAs and control RNAs.

This survey of sRNAs using the RiboGap database in Alphaproteobacteria, more importantly in Hyphomicrobiales, supports our hypothesis that *M. extorquens* most likely has more sRNAs than those already annotated.

The identification of sRNAs in bacteria is biased towards model organisms like *Escherichia coli* simply because they are more studied. We therefore decided to compare the size distribution of IGRs in *E. coli* K-12 substr. MG1655 (NC_000913.3) with those in *M. extorquens* AM1 (NC_012808.1) focusing on gaps between 50 and 1000 nucleotides (Figure 1, C-D). This specific bacterial strain was chosen since it is the sequenced strain the most closely related to the strain we used (ATCC55366). Seven distinct sRNAs are annotated in the genome of *M. extorquens* AM1 (ffh, ar45, CC2171, BjrC1505, BASRCI27, suhB and 5_ureB_sRNA). Small RNAs 5_ureB_sRNA, BASRCI27 and suhB are present more than once in the genome (in two, three and four copies respectively). Homology searches from Rfam covariance models from the entire sRNAs repertoire was performed on all available prokaryotic genomes in NCBI [^34^], leading to more sRNA predictions in RiboGap than in Rfam (Figure 1, C-D).

The size distribution of IGRs in the range of 50 and 1000 nucleotides for both proteobacteria has a similar pattern. In *E. coli*, approximately 65% of the annotated sRNAs are concentrated in the intergenic gaps between 50 to 400 nucleotides. Remarkably, only two trans-regulatory elements are annotated in this potential sRNA rich region in *M. extorquens*, reinforcing the idea that there are more to be discovered.

### Expression of annotated sRNAs

We first wanted to confirm the expression of some of the annotated sRNAs in *M. extorquens* by Northern blot analysis with bacteria grown with 1% methanol. Three of the annotated sRNAs in *M. extorquens* AM1 were selected to be experimentally validated (ffh, CC2171, BjrC1505) (Table 1). To use as positive controls for Northern blots, probes for 5S RNA, transfer-messenger RNA (tmRNA) and the leucine tRNA were created as well, all expected to be highly transcribed (Table 1). These would also act as a size guideline for the predicted sRNAs.

Hybridization was observed for all RNAs used as a positive control (5S RNA, tRNA-leu and tmRNA), confirming the proper transfer of the extracted RNA on the nitrocellulose membrane (Supplementary material, Figure S1). Bands were also detected for all three sRNAs that were annotated in the genome of *M. extorquens* (ffh, CC2171, BjrC1505), validating for the first time the presence of sRNAs in this biotechnologically relevant bacteria (Figure 2, A). All hybridization experiments were done in tri-replicates (data not shown).

**Figure 2.**
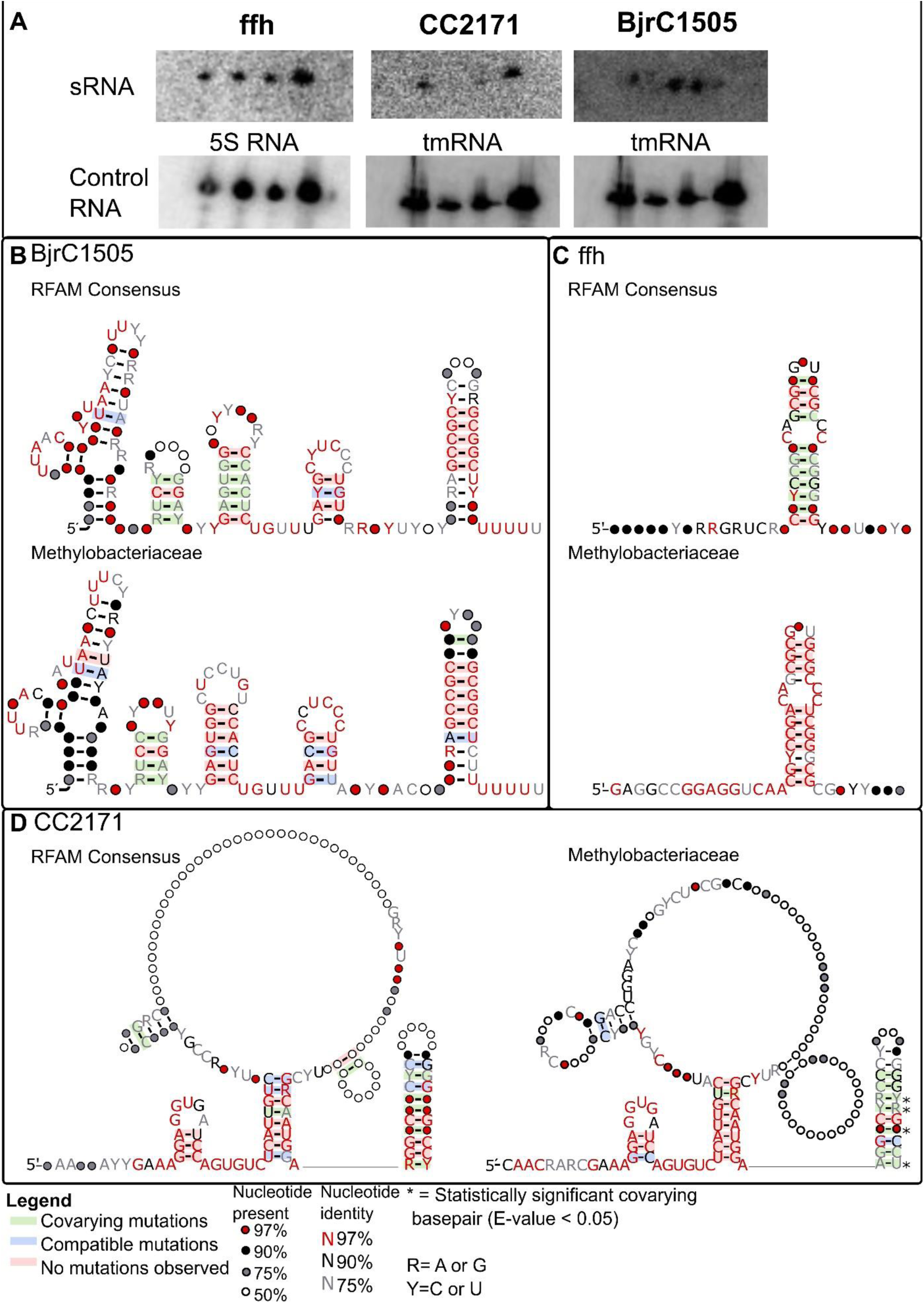
Structure of validated sRNAs by Northern Blot Analysis. (A) Hybridization of radioactively labeled probes for annotated sRNAs (ffh, CC2171 and BjrC1505). The four bands correspond to different time points (24h, 48h, 72h, 96h). For every membrane, a control RNA was also hybridized (5S RNA or tmRNA). The secondary structure for the sRNA BjrC1505, ffh, and CC2171 are depicted in (B), (C) and (D) respectively. For each RNA, the consensus structure from all bacteria (Rfam consensus) and that from Methylobacteriaceae are shown. Basepairs with statistically significant covariation are represented by * (E-value < 0.05) for Methylobacteriaceae secondary structure. Their significance was assessed with R-scape [^44^].

For every validated sRNAs, the secondary structure from the Methylobacteriaceae family corresponds to the Rfam consensus, which is the taxonomy classification of *M. extorquens*. However, more sequence conservation and less covariation are observed since the species are more closely related (Figure 2, B-C-D). Statistically significant covarying basepairs in Methylobacteriaceae can be observed only for the secondary structure of CC2171 (Figure 2, D). The sRNA CC2171 was first discovered within the bacteria *Caulobacter crescentus*, but the condition affecting this sRNA was not identified [^29^]. This is the first instance that the expression of sRNA CC2171 is validated in an Alphaproteobacteria other than *C. crescentus*. The sRNA ffh plays a role in the regulation of the *ffh* gene, which encodes for the cytoplasmic protein of the bacterial signal recognition particle (SRP) [^31^]. The cis-encoded sRNA found upstream of the gene ffh is well conserved and widespread amongst Alphaproteobacteria. Finally, the sRNA BjrC1505 was predicted to target acetolactate synthase, an enzyme important in the metabolism of branched-chained amino acids (BCAA) [^30, 35^]. This trans-activating sRNA is found in plant-associated Alphaproteobacteria such as in the family *Bradyrhizobiaceae* and some genera of the family *Rhizobiaceae*, but its expression had not been demonstrated by Northern Blot in another plant-associated bacteria, *M. extorquens* in this case.

### Prediction of sRNA Candidates

#### sRNA-Detect

Beyond these few, now confirmed, sRNAs, we were interested in discovering potential novel sRNAs. For this, we analyzed the transcriptome with sRNA-Detect [^23^]. This data came from a transcriptomic study realized as part of another research project (unpublished data). Briefly, *M. extorquens* strain ATCC55366 was genetically engineered to allow the accumulation of the tricarboxylic acid cycle (TCA) metabolite succinic acid using a ΔsdhA gap20::145 ΔphaC::Km^R^ triple mutant [^36^]. To investigate the impact of these mutations at the transcriptional level, RNA-seq data were acquired for the WT strain, the mutant at pH 6.5 and without pH control. We used the RNA-seq data from these three samples (each in triplicates) for the sRNA-Detect analysis, but without further focus on mutant vs WT strains, unless otherwise mentioned in the text.

Inspection of the transcriptome using sRNA-Detect resulted in a list of 10,267 detected candidates from all three conditions. Some of these were repetitive amongst growth conditions, with approximately 3,500 potential sRNAs for each of them. This includes multiples hits for a single ncRNA (e.g., 15 candidates are found within the 23S rRNA sequence). Sequences with a predicted length of 50 to 250 nucleotides by sRNA-Detect within the main chromosome (NC_012808.1) for the WT strain were kept for further analysis (2,079 candidates).

### Annotated RNAs among sRNA-Detect Candidates

Candidates were first inspected for the presence of already annotated RNA. The list of all annotated RNA within the genome of *M. extorquens* AM1 was obtained with RiboGap [^24^]. Amongst our list of presumptive regulatory elements, ribosomal RNAs (rRNA), sRNAs, transfer RNAs (tRNAs) and cis-regulatory elements were found (Table 2). Most importantly, we were able to recover six of the seven distinct sRNAs that are annotated in the genome of *M. extorquens* AM1 (all except 5_ureB_sRNA), confirming that sRNA-Detect is a reliable tool to detect sRNAs.

**Table 2.**
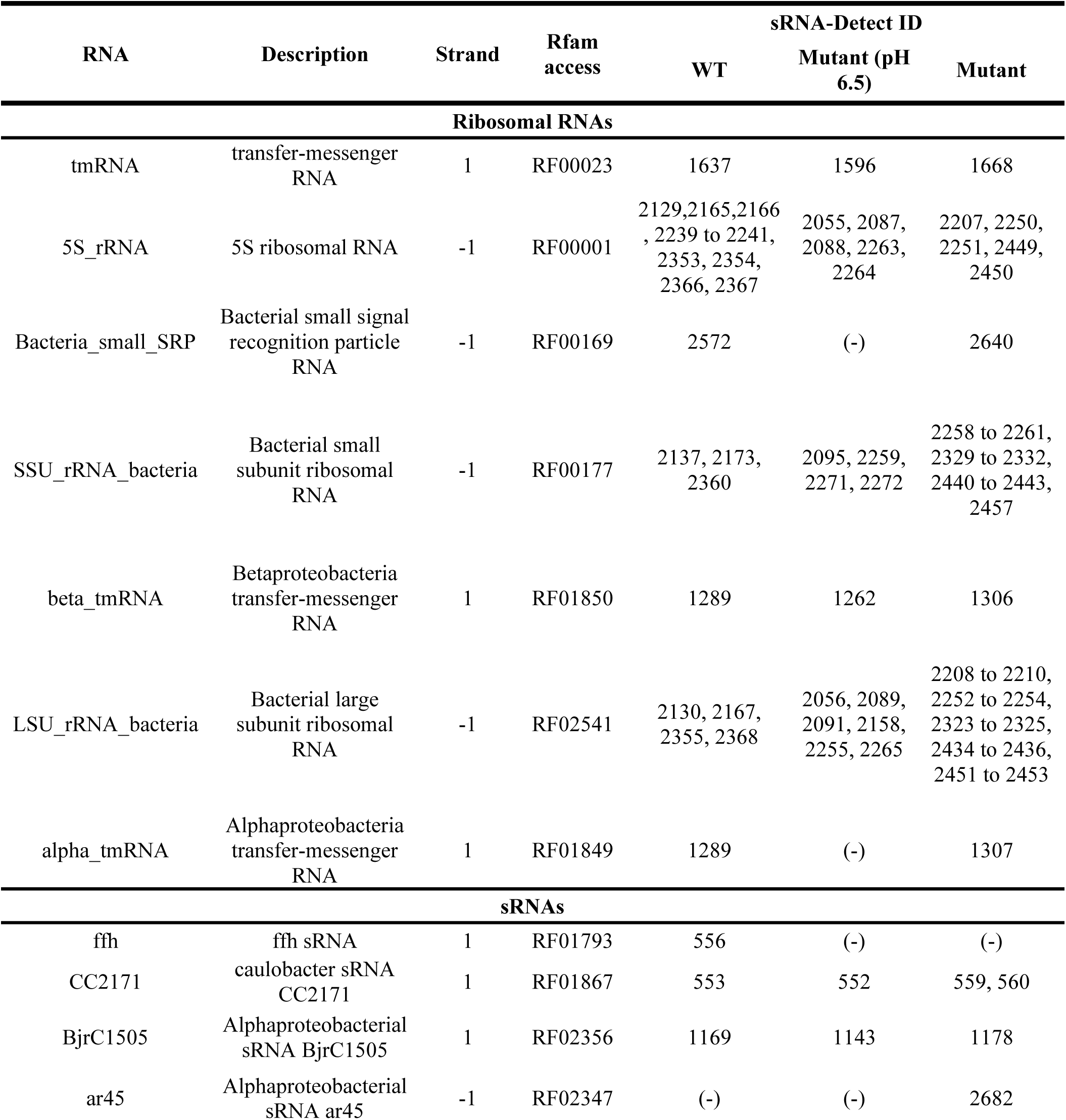
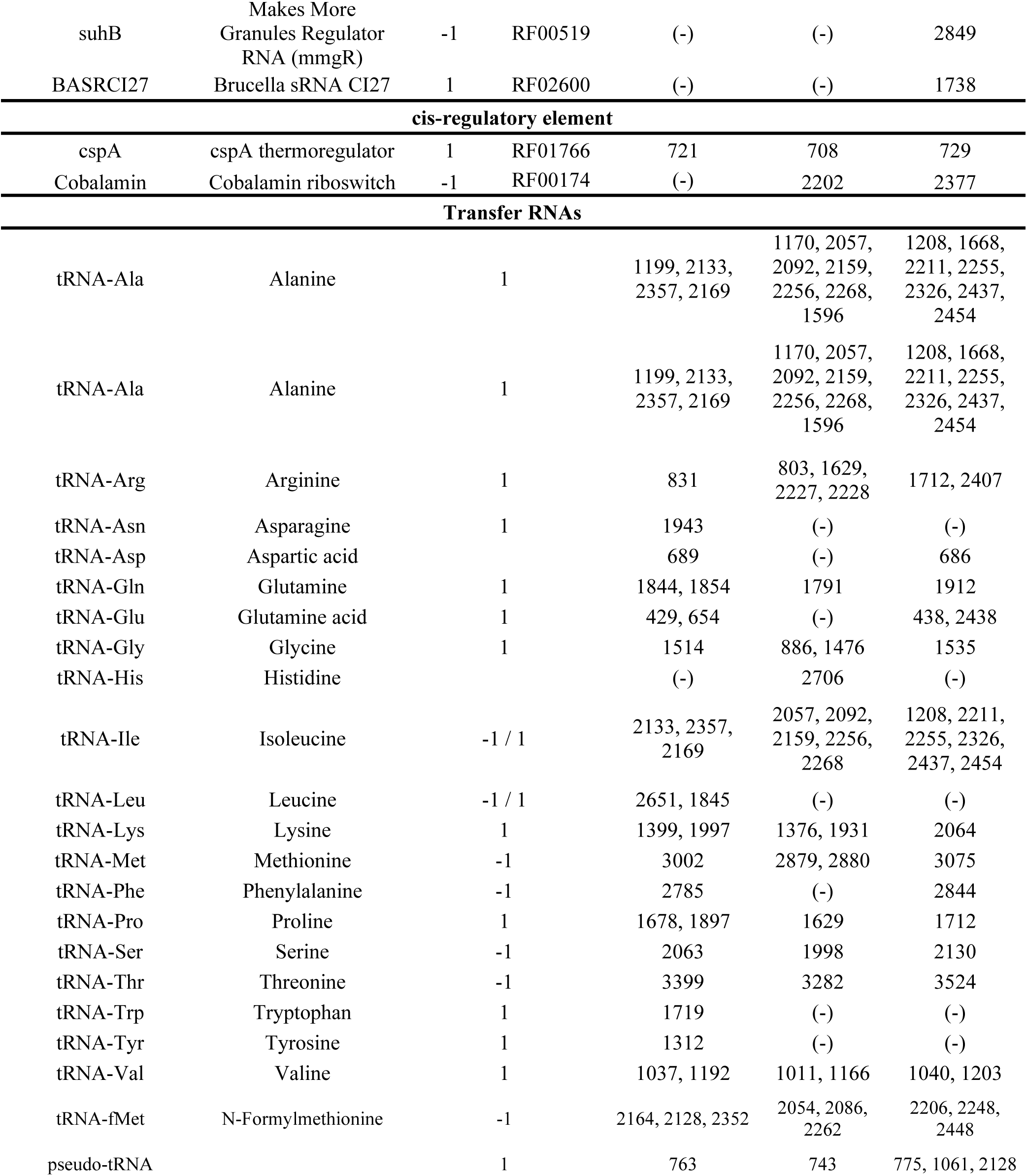
Annotated RNAs within sRNA-Detect candidates

Numerous candidates were proposed along the genome of *M. extorquens* AM1 for the WT strain. Amongst potential sRNA with a sRNA-Detect score higher than 1000 and a length between 50 to 250 nucleotides (388 candidates), 22 were selected to be tested experimentally (Supplementary material, Table S1). The value provided by sRNA-Detect represents the average read depth coverage, which is the sum of the reads mapped to each nucleotide of small transcripts divided by the length of such transcripts. It could therefore be interpreted as the level of expression of that RNA region. A cut-off of 1000 was determined to be acceptable since the mean score of all annotated RNAs was higher than this value (Figure 3).

**Figure 3.**
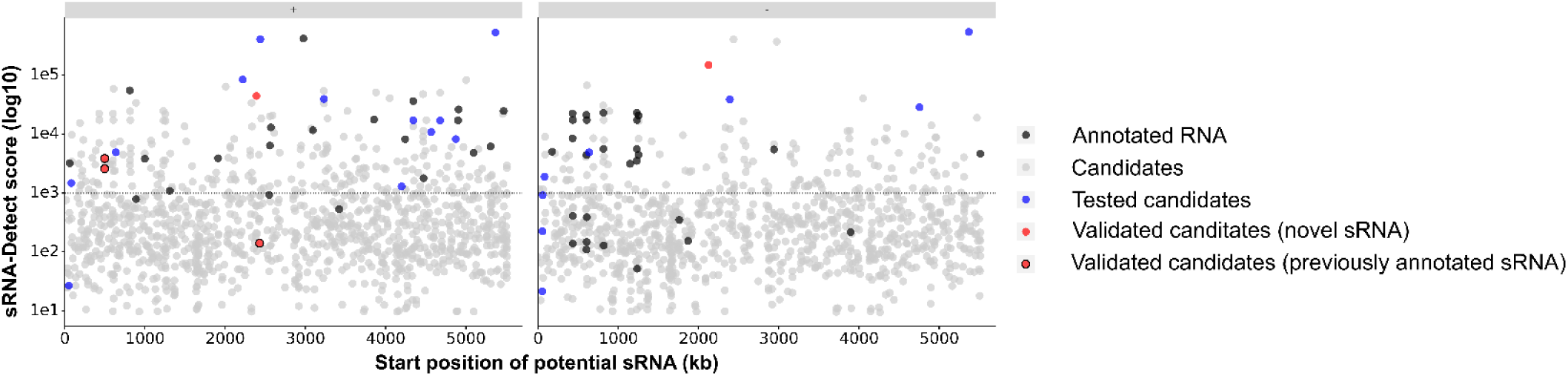
Putative sRNA candidates identified by sRNA-Detect. The x-axis represents the starting position of the putative RNA elements on the chromosome in kilobase pairs and the y-axis represents the score given by sRNA-Detect (log10). A cut-off of 1000 for this score is depicted by the dotted line. Already annotated RNAs with an E-value lower than 0.0005 are denoted in black. Blue dots illustrate candidates that could not be detected by Northern blot analysis. Red points show candidates where bands were observed with Northern blot analysis. Red dots with a black outline are sRNAs that were previously annotated, whereas red dots without an outline are novel sRNA specific to this study. Information is divided into positive (left panel) and negative strand (right panel).

Transcripts for sRNAs 1153 and 2624 were detected by Northern blot analysis (Figure 4). Hybridization of the probe for candidate 2624 was observed in triplicates with methanol or succinic acid as a source of carbon (data not shown). The band intensity for candidate 1153 was very weak, but could also be detected on two other membranes, albeit with an even weaker signal. When comparing their migration profile in a membrane with RNA of known size, both sRNA2624 and sRNA1153 are between 5S RNA (115 nt) and tmRNA (378 nt), where sRNA2624 is much closer to tmRNA. Further analysis focuses on sRNA2624, but a potential secondary structure was still determined for Met1153 based on IGR containing its sequence.

**Figure 4.**
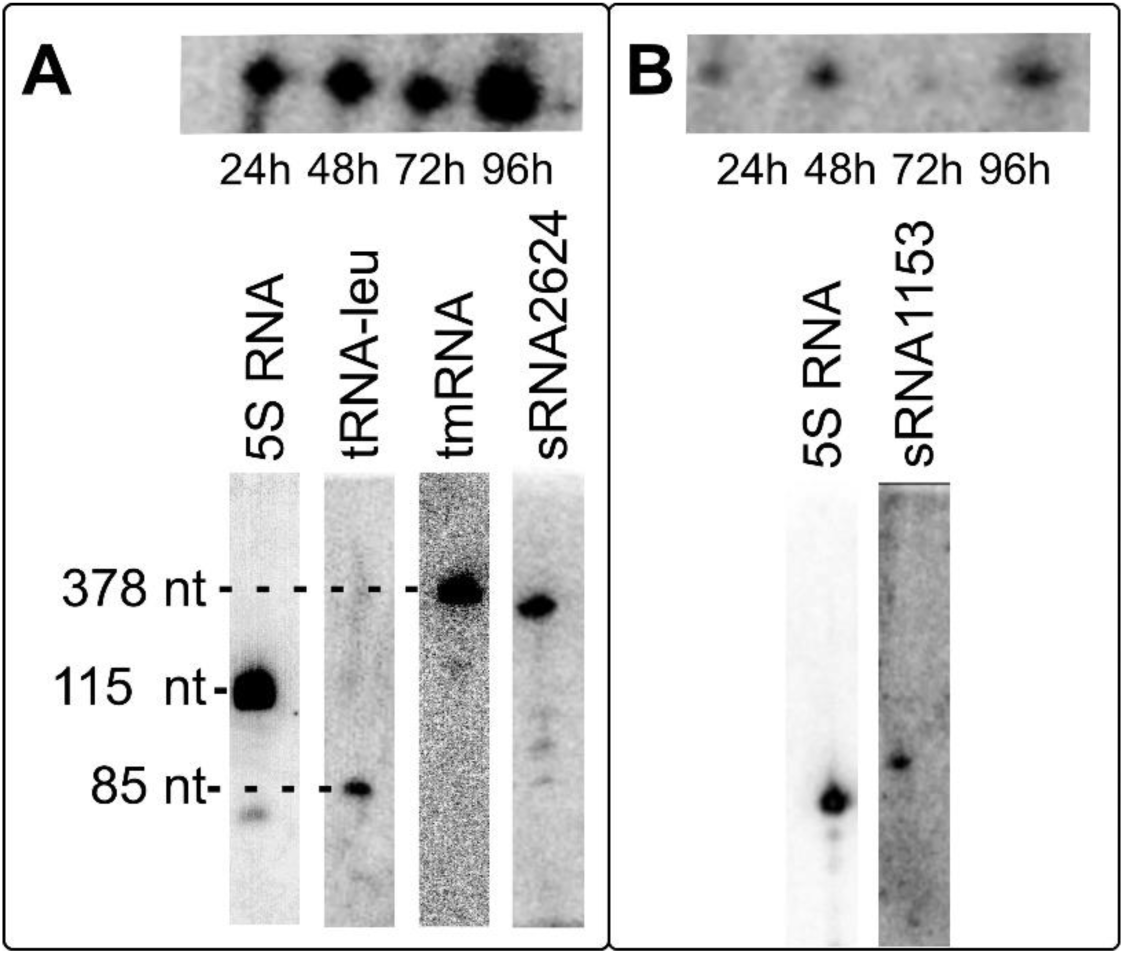
Validated expression of candidate sRNA2624 and sRNA1153 by Northern Blot analysis. (A) Hybridization of the probe complementary to candidate 2624 at different time points (24h, 48h, 72h and 96h). To estimate its size, probes for RNAs of known size were also hybridized on the same membrane (5S RNA, 115 nt; tRNA-leu, 85 nt and tmRNA, 378 nt). (B). Hybridization of the probe complementary to candidate 1153 at different time point (24h, 48h, 72h and 96h). To evaluate its size, a probe corresponding to 5S RNA (115 nt) was hybridized on the same membrane.

### Conserved genomic context of the candidate sRNA2624 in Methylobacteriaceae

The candidate sRNA2624 seems to be constitutively expressed over the growth of *M. extorquens* (Supplementary material, Figure S2). As controls, probes for the 5S RNA, leucine tRNA and tmRNA were also hybridized on the same membrane. By comparing their migration on the gel with that of the candidate sRNA2624, we can estimate its size to approximately 300 nucleotides (Supplementary material, Figure S1). According to sRNA-Detect prediction, sRNA2624 was only 105 nucleotides long and expressed within the negative strand. However, this bioinformatic method is known to miss some nucleotides at the 5’ and 3’ end, so it is probably longer, as suggested by the hybridization results (Figure 4) [^23^].

Using the BLASTn tool from NCBI [^37^], sequences for sRNA2624 and sRNA1153 were identified only in the genera *Methylorubrum* and *Methylobacterium* from the family Methylobacteriaceae (all intergenic regions containing sRNA2624 and sRNA1153 are listed in Supplementary material, Table S2 and S3 respectively). These regulatory elements were therefore named from Methylobacteriaceae and abbreviated, Met2624 and Met1153.

Within the bacteria encoding for Met2624, the genomic context varied. For 20 out of the 28 sequences with an annotated genomic context, the upstream gene encodes for an aspartate aminotransferase (Supplementary material, Figure S3, A-B). All instances of Met2624 were followed by a gene encoding a hypothetical protein of unknown function. The same protein of unknown function was encoded downstream of Met2624 for all bacteria recently reclassified within the new genus *Methylorubrum*, as well as a few *Methylobacterium* sp.; and a different gene was found downstream in other *Methylobacterium* species (Supplementary material, Figure S3). Met2624 was also compared to metagenomes within the NCBI GenBank [^38, 39^]. It was found in a biofilm metagenome coming from an environmental sample (PJQF01118147.1) and in ecological specimens taken from the Red Sea (FUFK012632339.1, FUFK010110014.1, FUFK010045813.1).

### Met2624: Small Protein or Functional RNA?

The program RNAcode [^40^] was ran to verify if Met2624 was a putative coding sequence. Small open reading frames (ORF) are often missed by gene annotations for proteins, posing a challenge to identify novel sRNAs. Software like RNAcode assess the coding potential of conserved regions to discriminate between protein coding and functional RNA. A multiple sequence alignment containing all IGRs where Met2624 is found, from both genomic and metagenomic data, was created with Clustal Omega [^41^] and submitted to RNAcode [^40^]. Results are represented in a list of hypothetical coding sequence, where their score and their P-value is provided (Supplementary material, Table S4). All identified potential coding sequence in a negative frame can be disregarded, since we know the transcribed strand coding for Met2624. RNAcode assigned a score to each presumed small ORF, where random noncoding regions generally do not have a scoring factor higher than 15 [^40^]. The results generated six potential ORFs in the (+) strand, with the sequence from metagenomic sample from *Methylobacterium nodulans* ORS 2060 as a reference for the position (the bacteria of reference is automatically selected by the program). Only one of those candidates had a scoring element higher than 15 (16.03) in the proper reading frame. It was associated with a P-value of 0.439 compared to the suggested limit of 0.01.

These results support our hypothesis that Met2624 did not encode for a small protein. To strengthen our claim, the secondary structure of Met2624 was analyzed with RNAz [^42^] to corroborate the idea that it is a functional RNA. The same multiple alignment file submitted to RNAcode was provided to the RNAz program, leading to a “RNA-class probability” of 0.94, indicating that it is most likely to be a functional RNA (Supplementary material, Figure S4). To make such forecast, RNAz considers the structure conservation index (SCI) and the thermodynamic stability (negative z-score). Functional RNAs are associated with a high SCI and thermodynamic stability, which were 0.66 and -2.03 respectively in the case of Met2624. A functional RNA is predicted when the probability is higher than 0.5, where high values correspond to more confident prediction (0.94 in the case of Met2624).

### Secondary Structure Predictions

Both RNAcode and RNAz results suggested that Met2624 is indeed a functional RNA and not simply an mRNA with a small open reading frame. All IGRs were aligned with Clustal Omega [^41^] to identify the more conserved regions containing Met2624 (sequences are underlined in Supplementary material, Table S2). A FASTA file containing all conserved regions from each intergenic sequences was submitted for secondary structure prediction with Graphclust [^43^] to establish a covariance model. Considering that Met2624 was only found in the family of Methylobacteriaceae, sequence conservation is relatively high, which limits covariation. Despite this conservation, some covariations and compatible mutations could be observed within the predicted conserved structure for Met2624 created (Figure 5). A potential secondary structure was also determined for Met1153 based on sequences containing the sRNA candidates in the same manner (Supplementary material, Figure S5).

**Figure 5.**
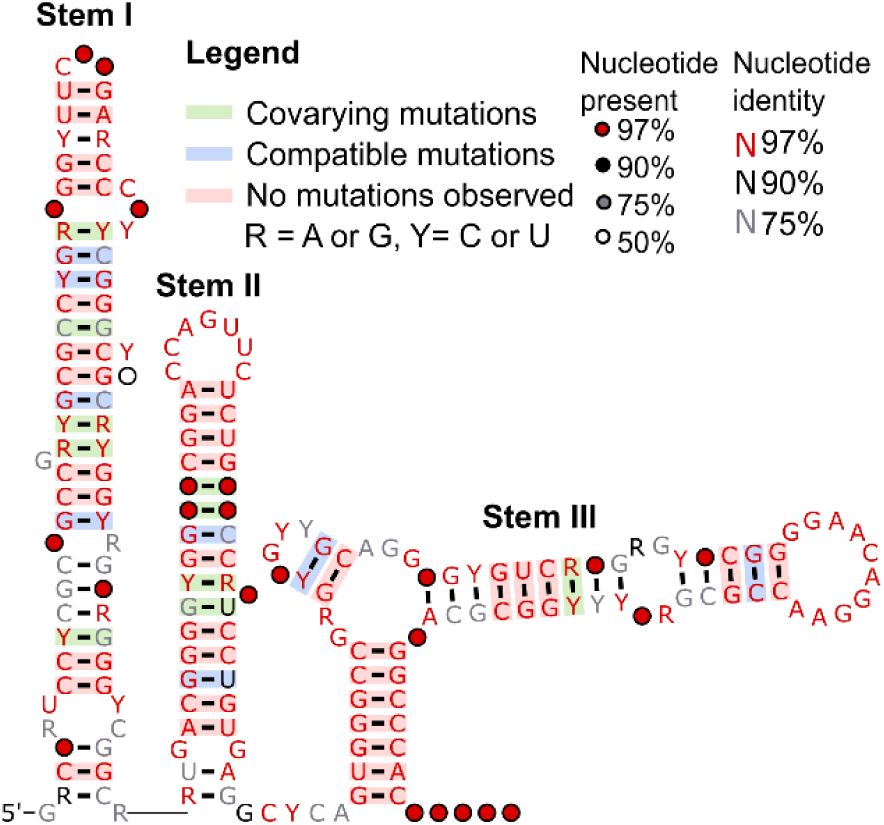
Secondary structure of Met2624. The structure was drawn by the program R2R [^53^]. Taken individually, none of the indicated covarying base pairs are considered statistically significant according to R-scape [^44^]. With alternatives Stockholm alignment, the base pair R-Y from stem III was covarying significantly based on R-scape (Supplementary material, Figure S6).

None of the covarying base pairs were statistically significant when evaluated with the R-scape tool for these predictions [^44^]. However, a base pair from stem III was covarying significantly based on R-scape for the structure of Met2624 when using an alternatives Stockholm alignment (Supplementary Material, Figure S5). Moreover, stems II and III are still predicted to form in this alternate conformation, even if the start and end positions of aligned sequences varied, further supporting our predicted secondary structures.

### Prediction of Promoter and Terminator for Met2624

The IGR from AM1 containing Met2624 was analyzed for the presence of a promoter and terminator (Figure 6). No promoters consistent with our Northern blot analyses were found with bTSSfinder [^45^]. This tool seeks classes of hypothetical promoters from *E. coli* (σ^70^ [RpoD/SigA], σ^38^ [RpoS], σ^32^ [RpoH] and σ^24^ [ECF subfamily]) [^45^]. These classes are all found in *M. extorquens*. Although it is considered as a complete program to predict promoters, it has a recall value in between 49% and 59% for known promoters [^45^] and it does not search for promoters associated with σ^54^, which are also found in *M. extorquens*. Hence, we turned to iPro54-PseKNC [^46^], to predict σ^54^ promoters [RpoN] (Figure 6, B).

**Figure 6.**
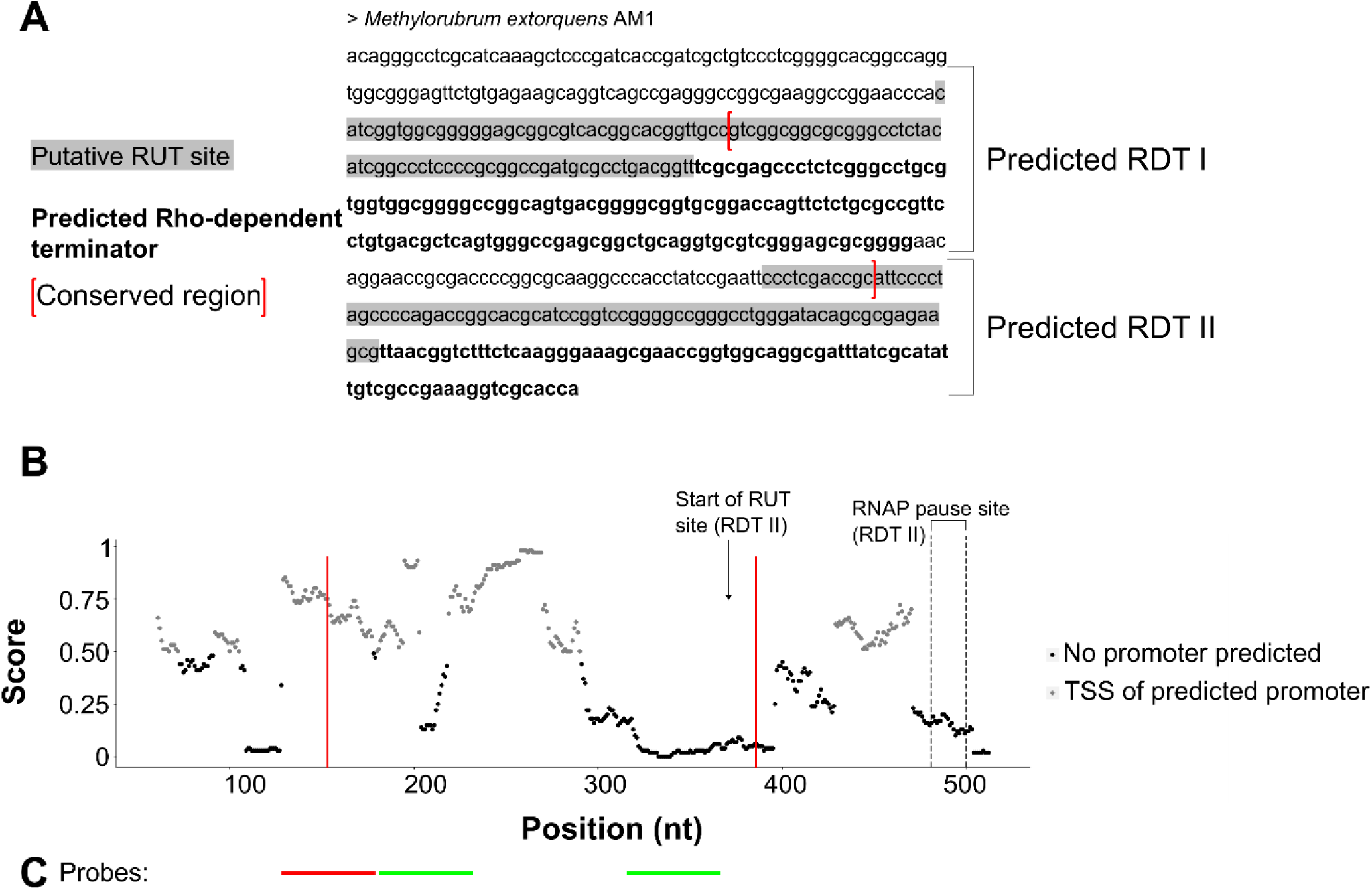
Promoter and terminator predictions for Met2624. (A) Sequence of intergenic region from *M. extorquens* AM1 containing Met2624. Two Rho-dependent terminators (RDT I and II) are predicted in this region by RhoTermPredict [^49^]. The minimum score given by the algorithm is 6, whereas the maximum score is 15 for predicted terminator. Their score was of 6 and 11 respectively. The Rho utilization sites (RUT) are highlighted in grey, whereas the region analyzed for RNA polymerase (RNAP) pause site are emphasized in bold. The start position of the RUT site and RNAP pause site of RDT II are respectively indicated in (B) by an arrow and delimited by two dashed lines. The intergenic region was analyzed for the presence of σ54 promoter with iPro54-PseKNC [^46^]. Every possible range of 81 nucleotides along the submitted sequence were analyzed. Dots represent the 61^st^ position of this range, where a TSS is expected if it is a promoter region. Grey dots represent the predicted TSS of regions containing a potential promoter, whereas black dots depict non promoter region. The score represents the probability to find a σ 54 promoter within a given range, where value higher than 0.5 are classified as putative promoter regions. The section delimited by the red brackets and lines in A and B corresponds to most conserved region when all IGR containing Met2624 are aligned. (C) Location of probes designed to detect Met2624. Probes in green lead to hybridization results corresponding to Met2624, whereas no hybridization was observed for the one in red, thus defining an approximate 5′ end of Met2624.

The program analyzed all possible 81 nucleotide ranges within the submitted sequence, and it identified different regions that could contain σ^54^ promoters. The transcription start site (TSS) of potential promoters are depicted in grey at the 61st position of the 81-nucleotide range. When validated with a set of known promoters, iPro54-PseKNC gave true TSS positions a score close to 1 [^46^]. Prokaryotic transcription is initiated by the recognition of the promoter sequence by sigma factors. They can be divided into two main categories defined by the type of sigma factors: σ^70^ and σ^54^. While σ^70^-dependent transcription controls the majority of housekeeping genes in normal condition, σ^54^ dependent transcription is extensively used by bacteria to regulate their metabolism in response to environmental changes [^46^]. Sigma 54 promoters are often associated with the transcription of genes related with carbon metabolism [^46^], making it an interesting promoter candidate for this C1 consumption model organism. Since sRNAs play an important regulatory role, it comes as no surprise that a σ^54^ promoter is predicted for Met2624.

No Rho-independent terminator was predicted with the tool RNIE in the IGR where Met2624 is found [^47^]. Met2624 regulation is therefore likely independent of the chaperone protein Hfq, since the poly-U tail of Rho-independent terminator is typically an important Hfq binding site [^48^]. Hfq can otherwise bind AU-rich regions, which are also absent from Met2624. However, two Rho-dependent terminators (RDT) were identified within the sequence using the tool RhoTermPredict [^49^] (Figure 6, A-B). The program assigns scores in between 6 and 15 to potential terminators, where the highest value represents the greater probability. RDT I and II had a value of 6 and 11 respectively.

To narrow down the location of a potential promoter and a terminator for Met2624, various probes towards the 5’ and 3’ were designed to delimit the sRNA experimentally (Figure 6, C). Predictions for promoters and terminators agree with our expected size following Northern blot analysis.

## Conclusion

This is the first time the expression of sRNAs was validated in the biotechnologically relevant bacterium *M. extorquens*. A better understanding of modes of regulations already in place within this bacterium could give hints on how to promote production yield. For example, recently implemented genetic tools like synthetic sRNAs [^50^] or CRISPR-Cas9 [51] could target newly discovered sRNAs that have an impact in the output of a desirable product or in the metabolism of a carbon source.

A list of potential sRNAs in *M. extorquens* was created through this research, which could be further analyzed to identify new and interesting candidates for future research. Most importantly, this study highlighted Met2624 as a sRNA specific to Methylobacteriaceae, even if we cannot completely exclude the possibility that it could encode a protein with a small ORF. In future research, the deletion or overexpression of this sRNA candidate could help elucidate its function and putative regulatory role. In any case, this research is the first to demonstrate the expression of sRNAs in the C1 metabolism model organism *M. extorquens*. It is meant to be the first overview of the role of sRNAs in *M. extorquens*, leading the way to future work including the characterization of Met2624 and the discovery of other novel sRNAs within *M. extorquens*, improving at the same time the value of *M. extorquens* as a biotechnological tool.

## Material and Methods

### Bioinformatics selection of candidates

#### RNA-sequencing data

A DASGIP® parallel bioreactor system (Eppendorf) equipped with 1.5 L reactor vessels was used to grow the *M. extorquens* wild-type strain ATCC55366 at pH 6.5 and its isogenic ΔsdhA gap20::145 ΔphaC::Km^R^ triple mutant at pH 6.5 and without pH control [^36^], each in biological triplicates, for a total of nine fermentation runs. Precultures were prepared as follow: two 3 L baffled Erlenmeyer containing 400 mL of CHOI Medium 4 [^10^], supplemented with 0.3% malic acid, were inoculated with cells harvested from freshly grown agar plates. Malic acid was supplemented to the growth culture to compensate for the ΔsdhA mutation that is interrupting the TCA cycle. Kanamycin was added to triple mutant precultures only (40 µg/mL). Precultures were incubated overnight at 30°C, under an agitation of 250 rpm. Then, precultures were used to seed reactors to obtain initial optical densities (600 nm) of approximately 0.25. Each reactor contained 1 L of CHOI Medium 4 supplemented with 0.3% malic acid. No antibiotic and no antifoam were used during fermentations. Air flow rate was set at 35 sL/h whereas dissolved oxygen was kept at 30% using solely an agitation cascade, as no pure oxygen supply was needed. The temperature was set 30°C and the pH was maintained using phosphoric acid (1M) and ammonium hydroxide (28%). Methanol concentration was also kept constant at 0.2% (v/v) using a methanol sensing and reading system (Intempco; Montréal, QC) coordinated with DASGIP’s feeding pumps. Commercial methanol was used (J.T. Baker®, HPLC grade). Reactors were run for 22-24 hours and achieved similar optical densities (2.7 ≤ x ≤ 3.4) (Supplementary Material, Table S5). Then, RNA extractions were performed using MasterPure^™^ RNA Purification Kit (Epicentre®) according to the manufacturer protocol. Absence of DNA contamination was confirmed by PCR. Furthermore, to testify RNA quality, all samples were submitted to Agilent 2100 Bioanalyser using Prokaryote Total RNA Nano Chips. Then, samples were sent to the Centre d’innovation Génome Québec et Université McGill (Montréal, QC) for the preparation of KAPA rRNA depleted libraries and Illumina® sequencing (HiSeq V4 – PE 125 pb sequencing lane).

#### sRNA-Detect

The input for the sRNA-Detect [^23^] is RNA-seq data aligned to a reference genome (SAM file). The genome of *M. extorquens* AM1 was taken from the NCBI database as a reference (NC_012808.1). The output is a list of potential sRNAs in a gene transfer format (GTF). To identify potential sRNAs, this method highlights RNA sequences that have a minimum depth coverage within a given range. Selected features also demonstrate low depth variation through their whole sequence. Already annotated regions are not classified as candidates. The complete method is described by Peña-Castillo *et al*. 2016 [^23^]. The sRNA-Detect workflow is accessible at http://www.cs.mun.ca/~lourdes/site/Welcome.html. A list of potential candidates was obtained for all three growth conditions of our RNA-seq experiment: WT, mutant and mutant with controlled pH. For all candidates, the following information was available: start and end positions, expected length, strand and sRNA-Detect score. As with any method, there is a risk of false positives, so it is important to consider other criteria. In this article for example, candidate Met2624 was further supported by the analysis of its genomic context and conservation, as well as predictions of terminator and promoter regions. Moreover, its coding potential was evaluated with bioinformatic programs like RNAcode [^40^] and RNAz [^42^] to support that it is a functional RNA.

#### RiboGap

This database can be accessed via a web server (www.ribogap.iaf.inrs.ca) [^24^]. It allows anyone to easily retrieve information on intergenic sequences of prokaryotes. Information on RNAs in RiboGap is extracted from the Rfam database [^25^]. Results are therefore limited to annotations within the Rfam database. A complete list of all annotated RNAs (including rRNAs, sRNAs and tRNAs) found in *M. extorquens* AM1 was created using RiboGap (Version 2). The RNA of interest was specified on the interface in the section ‘‘type’’ of RNA family, whereas the organism of concern was selected using their corresponding accession number. Only annotated RNAs with an E-value lower than 0.0005 were kept to be compared with sRNA-Detect candidates. This database was also used to retrieve all intergenic regions from *E. coli* K12 and *M. extorquens* AM1 to determine size distribution and presence of sRNAs. Finally, a list of all Alphaproteobacteria with annotated sRNAs was obtained from RiboGap.

### Bioinformatic Analysis of Candidates

Several bioinformatic tools were used to further characterize Met2624. Intergenic sequences containing the sequence for Met2624 and Met1153 were extracted from RiboGap [^24^] and the NCBI database [^34^], including metagenome sequences from GenBank [^38^] (Supplementary material, Table S2-S3). Information on proteins encoded upstream and downstream of Met2624 were also extracted from NCBI [^34^]. All sequences were aligned using the Clustal Omega interface (https://www.ebi.ac.uk/Tools/msa/clustalo/) [^41^] to identify a conserved region amongst them (underlined sequences in Supplementary material, Table S2).

To assess for the coding potential of our candidate sRNA, the conserved region containing all intergenic regions with Met2624 was submitted to RNAcode (version 0.3) [^40^]. The input of this program is an alignment file in the Clustal Omega format [^41^]. To validate it was a functional RNA, the same alignment file was submitted to RNAz [^42^], which is accessible as a tool in the Galaxy suite [^52^].

To predict a promoter region upstream of Met2624, the intergenic region from *M. extorquens* AM1 containing the candidate sRNA was analyzed with bTSSfinder [^45^]. The program takes as an input a FASTA file. Since no promoter in agreement with our experimental results was predicted, the interface iPro54-PseKNC was used to assess the presence of a σ^54^ promoter (http://lin-group.cn/server/iPro54-PseKNC) [^46^].

Information on the presence of a Rho-independent terminator was extracted from RiboGap [^24^]. Information on Rho-independent terminator on RiboGap comes from the RNIE program [^47^]. For Rho-dependent terminator predictions for Met2624, the intergenic regions from *M. extorquens* AM1 in a FASTA format was submitted to RhoTermPredict [^49^].

To predict the secondary structure of Met2624 and Met1153, a covariance model was created with the Graphclust [^43^] workflow on the Galaxy platform [^52^]. The consensus structure drawing was generated with R2R [^53^] using the alignment of the corresponding covariance model generated by Graphclust [^43^] with all intergenic regions containing the regulatory RNA. The alignment was generated with the cmalign tool from infernal [^54^]. For sRNAs CC2171, ffh, BjrC1505, the Rfam consensus structure was created using the Stockholm file from Rfam. The Methylobacteriaceae consensus was produced from the alignment of their covariance model with all sequences from this family containing the desired regulatory element from a FASTA file generated by RiboGap and aligned with cmalign. To evaluate whether the covarying basepairs were significant, the RNA multiple sequence alignment in a Stockholm format for each structure was submitted in R-scape (http://eddylab.org/R-scape/) [^44^].

Several figures in this article are represented with the help of ggplot2 [^55^] package within the Jupyter notebook [^56^].

The programs btSSfinder [^45^], RhoTermPredict [^49^], RNAcode [^40^], Infernal [^54^] and R2R [53] were ran in the Graham server from Compute Canada. RNAz [^42^] and Graphclust [^43^] were accessed via the Galaxy suite [^52^]. All other programs are accessible online via the provided links.

### Northern blot Analysis

#### Growth conditions of *M. extorquens*

Cultures of *M. extorquens* ATCC55366 were grown in 250 mL baffled Erlenmeyer flasks at 30 °C and 200 rotations per minutes (rpm) in CHOI Medium 4 as described in [^10^]. The CHOI growth medium corresponded to 1/5 of the volume of the baffled Erlenmeyer flask to allow for proper oxygenation. Methanol (1%) was added to the growth medium as a source of carbon and supplied every 24 hours (0.5%). The optical density at 600 nm was taken by spectrophotometry with Eppendorf BioSpectrometer ® for four days every 24 hours (Supplementary material, Figure S7). At every time point, 1 mL of culture was centrifuged at 5,000 rpm at 4 °C for 10 minutes and the supernatant was discarded. The bacterial pellet was stored at -80 °C before RNA extraction.

#### RNA Extraction

The bacterial pellets previously stored at -80 °C were lysed with 100 μL of a solution of 400 μg/mL of lysozyme in TE buffer (0.5 M EDTA and 1 M Tris-HCl, adjusted pH of 8.0) for 5 minutes at room temperature. RNA was extracted using TRIzol reagent (Invitrogen) as described in Rio *et al*., 2010 [^57^]. The RNA contained in the supernatant was precipitated by adding 2 volumes of 100% chilled ethanol and 0.1 volumes of 3 M sodium acetate (pH 5.2) and cooled at -80°C for at least 2 hours. The RNA was centrifuged at 4°C for 30 minutes at 14,000 rpm. The supernatant was discarded and 500 μL of 70% chilled ethanol was added to rinse the pellet. It was then centrifuged at 4°C for 5 minutes at 14,000 rpm. The supernatant was removed, and the pellet was left to dry for at least 15 minutes before being resuspended in RNAse-free water. The extracted RNA was quantified using a Nanodrop (Thermoscientific Nanodrop 2000).

#### Northern Blot

Ten micrograms of total RNA for each sample was migrated into a 10% denaturing 8 M urea polyacrylamide gel (PAGE) to separate the RNA according to its size. Samples were loaded with 2 X gel loading dye (0.05% bromophenol blue, 0.05% xylene cyanol, 10 mM EDTA [pH 8], 95% formamide) with TBE 1 X (0.09 M Tris-base, 1 mM EDTA, 0.09 M boric acid) as running buffer. The gel was migrated at 15 W for 1 hour. The RNA in the gel was transferred overnight to a nitrocellulose membrane with a positive charge (GE healthcare Amersham™ Hybond™ -N^+^) by capillary transfer using an assembly of Whatman® filter paper. The capillary transfer was set up as follows: 10X SSC (1.5 M sodium chloride and 0.15 M sodium citrate dihydrate) was poured into a container. Four Whatman® filter paper and a nitrocellulose membrane were cut to the length and width of the polyacrylamide gel. Another Whatman® filter paper was cut the same width, but its length was long enough to touch the buffer when placed on a support. They were all pre-soaked into 10X SSC buffer for 30 minutes prior to the assembly. The Whatman® filter papers, the nitrocellulose membrane and the polyacrylamide gel were all stacked one on top of the other.

The RNA was left to transfer from the polyacrylamide gel to the nitrocellulose membrane overnight. The next morning, the membrane was dried for a few minutes. To fix the RNA unto the membrane, shortwave UV light was used (UV stratalinker 2400 Stratagene). The membrane was stained with a methylene blue solution (0.02% methylene blue and 0.3 M sodium acetate pH 5.5) for 10 minutes with agitation to verify proper transfer of the RNA. The membrane was rinsed with distilled water for at least one hour. As the excess coloration was washed from the membrane, the bands corresponding to the highly abundant transferred RNA were revealed (data not shown).

### Hybridization of probes corresponding to candidate sRNAs

#### Radiolabelling of the DNA probe

For each candidate, a 50-nucleotide sequence complementary to the potential sRNA was selected in the middle of the sequence. Probes ordered from Integrated DNA technologies (IDT) were radiolabelled at the 5’ end with [γ-^32^P] ATP. A reaction was prepared with 0.5 µM of DNA probe, 1 X kinase buffer PNK, 20 µCi [γ-^32^P] ATP in a final volume of 20 uL. The labeling reaction was left to incubate for 1 hour at 37 °C. The labeled probes were purified on a 6 % denaturing gel (8 M urea PAGE, polyacrylamide gel electrophoresis). Loading dye 2 X and 1 X TBE was used as described before. The gel was exposed with phosphor imaging screens for 5 minutes before being scanned with a Typhoon^™^ FLA9500 (GE Healthcare Life Sciences). The bands corresponding to the probes were cut out of the gel and conserved at -20°C for future work. Intensity of radioactive bands was quantified with ImageJ.

The nitrocellulose membrane with the transferred RNA was pre-incubated with 15 mL hybridization buffer (20 X SSC, 50 X Denharts solution [2% Bovine Serum Albumin (BSA), 2% Ficoll 400, 2% Polyvinylpyrrolidone (PVP)], 10% sodium dodecyl sulfate [SDS], 100 µg/mL salmon sperm [Invitrogen, Thermofisher scientific]) in a rotating hybridization oven at 42 °C for 1 hour in a flask. After pre-incubation, the gel fragment containing a radiolabelled DNA probe was added inside the flask and left to incubate overnight in a rotating oven. The next day, the gel fragment and the hybridization buffer were recovered from the flasks and stored at -20°C for future work. The same probe can be used for many experiments if the DNA probe is still radioactive. The nitrocellulose membrane was washed in four steps as follows: 1 minute with washing solution I (2 X SSC buffer, 0.1 % SDS), 5 minutes with washing solution I, and twice for 10 minutes with washing solution II (0.2 X SSC buffer, 0.1 % SDS). All washing steps were performed in the rotation oven at 42 °C. The washed membranes were wrapped into Saran plastic wrap and exposed on phosphor imaging screens overnight before being scanned by a Typhoon^™^ FLA9500 (GE Healthcare Life Sciences). Intensity of radioactive bands were quantified with ImageJ. Membranes containing RNA can be used several times with different probes, if they are washed between each candidate with washing solution III (0.1X SSC buffer, 0.1% SDS) for 2 hours at 80 °C. To ensure that the membranes were cleaned, they were exposed in phosphor imaging screens as before. If radioactivity was still present, the last washing step was repeated. The membrane was stored in Saran wrap plastic between uses.

## Data availability statement

Data available on request from the authors

## Authors Contributions

EB carried and conceived most of the experiments and wrote the manuscript. SD and KT helped with the bioinformatic analysis. MJL and MW helped with set-up of fermentation experiments for RNA-seq. MGL performed RNA extractions for RNA-seq. RI, MGL and CBM contributed with the experiment design and initiated the research project. JP supervised the project and revised the manuscript.

## Acknowledgements

The authors would like to thank Jessie Muir for the assistance with writing R code and the laboratory of Charles Greer for assistance and sharing of equipment. The authors would also like to acknowledge Aurélie Devinck, Quetia Joseph and Philip Loranger for the revision of this manuscript. This research was enabled in part by support provided by Calcul Québec (www.calculquebec.ca) and Compute Canada (www.computecanada.ca).

