## Supplementary material for "Analysis of non-coding RNAs in *Methylorubrum extorquens* reveals a novel small RNA specific to Methylobacteriaceae"

### Analysis of non-coding RNAs in Methylorubrum extorquens reveals a novel small RNA specific to Methylobacteriaceae

Supplementary material

#### Tables

##### Table S1. Probes for candidates tested by Northern blot analysis

| ID | Start | End | Size | Strand | Probe |
|---|---|---|---|---|---|
| 426 | 51119 | 51301 | 182 | + | GGCAGCCCAGCCTGCTGCTCCTTCAGGATGCCGATGATCTGCTCTTCGCT |
| 432 | 80737 | 80895 | 158 | + | CGGGCTCACGGTCTTTGGCCGTGAACCCTAAAAGTGCGGCGATATGAACC |
| 609 | 636257 | 636362 | 105 | + | GAAGGATCCTCCAGGGATCGCGCGGGATCTGGACGATCCTTCGTGGCCGC |
| 1113 | 2218259 | 2218369 | 110 | + | TGTGCTGATGTTTCCGCTTCTGCGGATCACCGTCACGAAAAACCCGAGCGG |
| 1153 | 2388747 | 2388799 | 52 | + | TCTGGAGCCCTCCTTCGAGGCCTCCGCTTCGCTCCGGCACCTCAGGATGA |
| 1175 | 2436125 | 2436237 | 112 | + | AGCTCGTCAGGCTCATAACCTGAAGGTCGCTGGTTCAAATCCAGCCCCCG |
| 1348 | 3229332 | 3229428 | 96 | + | AAGCTCGGCCGTATGTGAACCTTGGCCGTTCGGCAGCGTTTCCCGAATGG |
| 1627 | 4198109 | 4198315 | 206 | + | CACGCGGGGAGAGGGTCGCGACGACCACGG |
| 1679 | 4343857 | 4343949 | 92 | + | CGGAACTTGCAAACCATCTGCAAGCCGTAACCCCGACGCCGAAGGCATCA |
| 1752 | 4567059 | 4567168 | 109 | + | CGCGACGACCGGATGCGCCCTCGACGGGCGCTTCCGCCGCCGGCACGATC |
| 1776 | 4678233 | 4678297 | 64 | + | GCCTGCGAGCGCCCTGCCGGGCACTTCACGTGGGCCATCCGCCGGAACGG |
| 1819 | 4874156 | 4874225 | 69 | + | GTGGGATCGCGTTGGCCGCTATCCCGCGCT |
| 1969 | 5367912 | 5368067 | 155 | + | CGTCCGGTGTTCGTGGCGGGCCACTCAGTCCAAGACCGAACGGGTTCTCA |
| 2036 | 51121 | 51275 | 154 | - | AGACGGACGATGAAGAAGAGCCGGTTTAGCGAAGAGCAGATCATCGGCAT |
| 2038 | 54339 | 54419 | 80 | - | TCTGATCGTTGCCTCGCGTGCTAGACCCATGCGCGGGTCGCCGCAATGCG |
| 2039 | 57158 | 57258 | 100 | - | TAGCGCCGCCCGAGGCGCGCGGGCAAGAGCCCGTTCCGGCCACGCCGGTA |
| 2043 | 80757 | 80895 | 138 | - | AATGCCTTCGGGCTCACGGTCTTTGGCCGTGAACCCTAAAAGTGCGGCGA |
| 2185 | 636269 | 636362 | 93 | - | AGCGGCCACGAAGGATCGTCCAGATCCCGCGCGATCCCTGGAGGATCCTTC |
| 2624 | 2125529 | 2125634 | 105 | - | AACGGCGCAGAGAACTGGTCCGCACCGCCCCGTCACTGCCGGCCCCGCCA |
| 2674 | 2388747 | 2388799 | 52 | - | CCTCATCCTGAGGTGCCGGAGCGAAGCGGAGGCCTCGAAGGAGGGCTCCA |
| 3211 | 4753719 | 4753781 | 62 | - | GACACCTTCAACTAGAAGGCGTCGCCGGACTCGATCCGGTGAGAACG GG |
| 3355 | 5367928 | 5368087 | 159 | - | TGAGAACCCGTTCGGTCTTGGACTGAGTGGCCCGCCACGAACACCGGACG |

#### Table S2. Intergenic regions containing Met2624.

More conserved region containing Met2624 based on alignment results of all intergenic regions are italicized.

| **Hits from Genomic data** |
|---|
| >*Methylorubrum extorquens* strain TK0001 |
| ACAGGGCCTCGCATCAAAGCTCCCGATCACCGATCGCTGTCCCTCGGGGCACGGCCGCGTGGCGGGA |
| GTTCTGTGAGAAGCAGGTCAGCCGAGGATCGGGAAGGGCCGGAACCCACATCGGTGGCAGGGGAGC |
| GGCGTCACGGCACGGTTGCCGTCGGCGGCGCGGGCCTCTACATCGGCCCTCCCCGCGGCCGATGCGC |
| CTGACGGTTTCGCGAGCCCTCTCGGGCCTGCGTGGTGGCGGGGCCGGCAGTGACGGGGCGGTGCGG |
| ACCAGTTCTCTGCGCCGTTCCTGTGACGCTCAGTGGGCCGAGCGGCTGCAGGTGCGTCGGGAGCGCG |
| GGGAACAGGAACCGCGACCCCGGCGCAAGGCCCACCTATCCGAATTCCCTCGACCGAATTCCCCTAG |
| CCCCTGGCCGGCACGCATCCGGTCCGGGGCCGGGCCTGGGATACAGCGCGAGAAGCGTTAACGGTC |
| TTTCTCAAGGGAAAGCGAACCGGTAGCAGGCGATTTATCGCAGATTGTCGCCGAAAGGTCGCACCA |
| >*Methylorubrum zatmanii* strain PSBB041 |
| ACAGGGCCTCGCATCAAAGCTCCCGATCACCGATCGCTGTCCCTCGGGGCACGGCCGCGTGGCGGGA |
| GTTCTGTGAGAAGCAGGTCAGCCGAGGATCGGGAAGGGCCGGAACCCACATCGGTGGCAGGGGAGC |
| GGCGTCACGGCACGGTTGCCGTCGGCGGCGCGGGCCTCTACATCGGCCCTCCCCGCGGCCGATGCGC |
| CTGACGGTTTCGCGAGCCCTCTCGGGCCTGCGTGGTGGCGGGGCCGGCAGTGACGGGGCGGTGCGG |
| ACCAGTTCTCTGCGCCGTTCCTGTGACGCTCAGTGGGCCGAGCGGCTGCAGGTGCGTCGGGAGCGCG |
| GGGAACAGGAACCGCGACCCCGGCGCAAGGCCCACCTATCCGAATTCCCTCGACCGAATTCCCCTAG |
| CCCCTGGCCGGCACGCATCCGGTCCGGGGCCGGGCCTGGGATACAGCGCGAGAAGCGTTAACGGTC |
| TTTCTCAAGGGAAAGCGAACCGGTAGCAGGCGATTTATCGCAGATTGTCGCCGAAAGGTCGCACCA |
| >*Methylorubrum extorquens* strain PSBB040 |
| ACAGGGCCTCGCATCAAAGCTCCCGATCATCGATCGCTGTCCCTCGGGGCACGGCCGCGTGGCGGGA |
| GTTCTGTGAGAAGCAGGTCAGCCGAGGACCGGCGAGGGCTAGAACCCATGTCGGCGGCGGGGGAGC |
| GGCGTCACGGCACGGTTGCCGTCGGCGGCGCGGGCCTCTACATCGGCCCTCCCCGCGGCCGATGCGC |
| CTGACGGTTTCGCGAGCCCTCTCGGGCCTGCGTGGTGGCGGGGCCGGCAGTGACGGGGCGGTGCGG |
| ACCAGTTCTCTGCGCCGTTCCTGTGACGCTCAGTGGGCCGAGCGGCTGCAGGTGCGTCGGGAGCGCG |
| GGGAACAGGAACCGCGACCCCGGCGCAAGGCCCACCTATCCGAATTCCCTCGACCGCATTCCCTTAG |
| CCCCTGGCCGGCACGCATCCGGTCCGGGGCCGGGCGTGGGATACAGCGCGAGAAGCGTTAACGGTC |
| TTTCTCAAGGGAAAGCGAACCGGTGGCAGGCGATTTATCGCAGATTGTCGCCGAAAGGTAGCACCA |
| >*Methylobacterium* sp. AMS5 |
| GGCGGTCCTTCGGGGCAAAGCGTCGCCCGACGGGCAGCGGCGTCACGGCACGGTTGCCGTCGGCGG |
| CGCAGGCCTCTACATCGGCCCTCCCCGCGGCCGATGCGCCCGATGGTTTCGCGAGCCCTCTCGGGCC |
| TGCGTGGTGGCGGGGCCGGCAGTGACGGGGCGGTGCGGACCAGTTCTCTGCGCCGTTCCTGTGACGC |
| TCAGTGGGCCGAGCGGCTGCAGGTGCGTCGGGAGCGCGGGGAACAGGAACCGCGACCCCGGCGCAA |
| GGCCCACCTACCTGAATTCCCTCGCCCGAATTCCCTTGGCCCCAGACCGGCACGCGTCCGGTCCGGG |
| GCCGGGCCTGGGATATCGGCGAGCGAAGCGTTAACGGTCTTTCTCAAGGGAAAGCGAACCGCTCGT |
| GGGCGATCTATCGCAGACCATCGCCTAAAGGTCGCACCA |
| >*Methylorubrum extorquens* strain DM4 |
| ACACGGCCTCGCATCAAAGCTCCCGATCACCGATCGCTGTCCCTCGGGGCACGGCCGCGTGGCGGGA |
| GTTCTGTGAGAAGCAGGTCAGCCGAGGGCCGGCGAAGGCTGGAACGTACATCGGTGGCGGGGGAGC |
| GGCGTCACGGCACGGTTGCCGTCGGCGGCGCGGGCCTCTACATCGGCCCTCCCCGCGGCCGATGCGC |
| CTGACGGTTTCGCGAGCCCTCTCGGGCCTGCGTGGTGGCGGGGCCGGCAGTGACGGGGCGGTGCGG |
| ACCAGTTCTCTGCGCCGTTCCTGTGACGCTCAGTGGGCCGAGCGGCTGCAGGTGCGTCGGGAGCGCG |
| GGGAACAGGAACCGCGACCCCGGCGCAAGGCCCACCTATCCGAATTCCCTCGACCGCATTCCCTAAG |
| CCCCAGACCGGCACGCATCCGGTCCGGGGCCGGGCCTGGGTACAGCGCGAGAAGCGTTAACGGTCT |
| TTCTCAAGGGAAAGCGAACCGGTGGCAGGCGATTTATCGCAGATTGTCGCCGAAAGGTCGCACCA |
| >*Methylorubrum extorquens* AM1 |
| ACAGGGCCTCGCATCAAAGCTCCCGATCACCGATCGCTGTCCCTCGGGGCACGGCCAGGTGGCGGG |
| AGTTCTGTGAGAAGCAGGTCAGCCGAGGGCCGGCGAAGGCCGGAACCCACATCGGTGGCGGGGGAG |
| CGGCGTCACGGCACGGTTGCCGTCGGCGGCGCGGGCCTCTACATCGGCCCTCCCCGCGGCCGATGCG |
| CCTGACGGTTTCGCGAGCCCTCTCGGGCCTGCGTGGTGGCGGGGCCGGCAGTGACGGGGCGGTGCG |
| GACCAGTTCTCTGCGCCGTTCCTGTGACGCTCAGTGGGCCGAGCGGCTGCAGGTGCGTCGGGAGCGC |
| GGGGAACAGGAACCGCGACCCCGGCGCAAGGCCCACCTATCCGAATTCCCTCGACCGCATTCCCCTA |
| GCCCCAGACCGGCACGCATCCGGTCCGGGGCCGGGCCTGGGATACAGCGCGAGAAGCGTTAACGGT |
| CTTTCTCAAGGGAAAGCGAACCGGTGGCAGGCGATTTATCGCATATTGTCGCCGAAAGGTCGCACCA |
| >*Methylorubrum extorquens* CM4 |
| ACAGGGCCTCACAACAAGGCTCCCGATCATCGATCGCTGTCCCTCGGGGCACGGCCGCGTGGCGGG |
| AGTTCTGTGAGAAGCAGGTCAACCGAGGACCGGCGAGGGCCAGACCCCACGTCGGCGGCGGGGGAG |
| CGGCGTCACGGCACGGTTGCCGTCGGCGGCGCGGGCCTCTACATCGGCCCTCCCCGCGGCCGATGCG |
| CCTGACGGTTTCGCGAGCCCTCTCGGGCCTGCGTGGTGGCGGGGCCGGCAGTGACGGGGCGGTGCG |
| GACCAGTTCTCTGCGCCGTTCCTGTGACGCTCAGTGGGCCGAGCGGCTGCAGGTGCGTCGGGAGCGC |
| GGGGAACAGGAACCGCGACCCCGGCGCAAGGCCCACCTATCCGAATTCCCCTAGCCCCTGGCCAGC |

Supplementary material

ATGCATCCGGTCCGGGGCCGGGCCTGGGATACAGCGCGAGAAGCGTTAACGGTCTTTCTCAAGGGA
AAGCGAACCGGTGGCAGGCGATTTATCGCAGATTGTCGCCGAAAGGTCGCACCA
>*Methylobacterium* sp. NI91
AGCGGCACGTCGGCCTTGCGTCGCCGGAAGGGCAGCGGCGTCACGGCGCGGTTGCCCGCGGCGGCG
CGGACCTCTACATCGGCCCTCCCCGCGGCCGATGCGCCTGATGGTTTCGCGAGCCCTCTCGGGCCTG
CGTGGCGGCGGGGCCGGCAGTGACGGGGCGGTGCGGACCAGTTCTCTGCGCCGTTCCTGTGACGCTC
AGTGGGCCGAGCGGCTGCAGGTGTGTCGGAAGCGCGGGGAACAGGAACCGCGATTTCGGCGCATGG
CCCACCCGATTCCCAGCCTTGCCGATCCGGGCTCCAGCCCGGATCGGGAACGGGTTCCTGGGATGGC
GCCAGAGCCGTTAACGGTCTTTCGCAAGGGAAAACCAACCAGACGCATGCAACCATTCTGCGCTTTG
TCTCAGAAAGGTTGCACAA
>*Methylobacterium* sp. CLZ
AGCGGCACGTCGGCCTTGCGTCGCCGGAAGGGCAGCGGCGTCACGGCGCGGTTGCCCGCGGCGGCG
CGGACCTCTACATCGGCCCTCCCCGCGGCCGATGCGCCTGATGGTTTCGCGAGCCCTCTCGGGCCTG
CGTGGCGGCGGGGCCGGCAGTGACGGGGCGGTGCGGACCAGTTCTCTGCGCCGTTCCTGTGACGCTC
AGTGGGCCGAGCGGCTGCAGGTGTGTCGGAAGCGCGGGGAACAGGAACCGCGATTTCGGCGCATGG
CCCACCCGATTCCCAGCCTTGCCGATCCGGGCTCCAGCCCGGATCGGGAACGGGTTCCTGGGATGGC
GCCAGAGCCGTTAACGGTCTTTCGCAAGGGAAAACCAACCAGACGCATGCAACCATTCTGCGCTTTG
TCTCAGAAAGGTTGCACAA
>*Methylorubrum populi* strain YC-XJ1
CTGGAAGAGGCCTGCCGCCGCATCCAGCGCTTCTGCGGCTCGCTGCGGTAGGCGGCGCCACGGGGC
AAAGCGTCGCCGAAGGGGCAGCGGCGTCACGGCGCGGTTGCCCGCGGCGGCACAGACCTCTACATC
GGCCCTCCCCGCGGCCGATGCGCCTGACGGTTTCGCGAGCCCTCTCGGGCCTGCGTGGCGGCGGGGC
CGGCAGTGACGGGGCGGTGCGGACCAGTTCTCTGCGCCGTTCCTGTGACGCTCAGTGGGCCGAGCGG
CTGCAGGTGCGTCGGGAGCGCGGGGAACAGGAACCGCGACCCCGGCGCAAGGCCCACACGATTCCT
GCGCTCTCCGATCCGGCCTCGAGCCCGGATCGGGGGCTGCCTTCCGGGCGCCGCGCGAGAACCGTTA
AGGGTCTTTCGCAAGGGAAAACAAACCGGCTCCATGCAACCTTTCGCACGTTGTCTCGAAAAGGTTG
CACAA
>*Methylobacterium* sp. DM1
AGCCGCACGTCGGCCTTGCGTCGCCGGAAGGGCAGCGGCGTCACGGCGCGGTTGCTCGCGGCGGCG
CGGACCTCTACATCGGCCCTCCCCGCGGCCGATGCGCCTGATGGTTTCGCGAGCCCTCTCGGGCCTG
CGTGGCGGCGGGGCCGGCAGTGACGGGGCGGTGCGGACCAGTTCTCTGCGCCGTTCCTGTGACGCTC
AGTGGGCCGAGCGGCTGCAGGTGTGTCGGAAGCGCGGGGAACAGGAACTGCGATTTCGGCGCATGG
CCCACCCGATTCCCAGCCTCGCCGATCCGGGCTCCAGCCCGGATCGGGAACGGTGTTCCCGGGTTGG
CGCCAGATCCGTTAACGGTCTTTCGCAAGGGAAAACCAACCAGACGCATGCAACCATTTTGCGCTTT
GTCTCAGAAAGGTTGCACAA
>*Methylorubrum populi*
TGCCATCCAGACTAGCAGGTTCGATCCCGGCTTTAACGCCGCGGGGCACGGCCGGCCGATCCCGCGA
TGCGCAGAGCGCATCGCGGCGTCACGGCGCGGTTGCCCGCGGCGGCGCGGGCCTCTACATCGGCCCT
CCCCGCGGCCGATGCGCCTGACGGTTTCGCGAGCCCTCTCGGGCCTGCGTGGCGGCGGGGCCGGCAG
TGACGGGGCGGTGCGGACCAGTTCTCTGCGCCGTTCCTGTGACGCTCAGTGGGCCGAGCGGCTGCCG
GTGCGTCGGGAGCGCGGGGAACAGGAACCGCGATCCCGGCGCAAGGCCCACCTCTCTGCCTCCCCA
TCACAGGCCGAACAGCACCGGTCCTCGCACGCCGAACCGGAGCGCCGTTAACGGTCTTTCTCAAGGG
AAAACAAACCGCTCGCAGGCAATCTTTCGCACGTTGTCCGAGAAAGGTCGCTCCA
>*Methylorubrum populi* BJ001
GGCGGCGCCACGGGGCAAAGCGTCGCCGAAGGGGCAGCGGCGTCACGGCGCGGTTGCCCGCGGCGG
CACGGACCTCTACATCGGCCCTCCCCGCGGCCGATGCGCCTGACGGTTTCGCGAGCCCTCTCGGGCC
TGCGTGGCGGCGGGGCCGGCAGTGACGGGGCGGTGCGGACCAGTTCTCTGCGCCGTTCCTGTGACGC
TCAGTGGGCCGAGCGGCTGCAGGTGCGTCGGGAGCGCGGGGAACAGGAACCGCGACCCCGGCGCAA
GGCCCACACGATTCCTGTGCTCTCCGATCCGGCCTTAAGCCCGGATCGGGGGCTGCCTTCCGGGCGC
CGCGCGAGAACCGTTAAGGGTCTTTCGCAAGGGAAAACAAACCGGCTCCATGCAACCTTTCGCACGT
TGTCTCGAAAAGGTTGCACAA
>*Methylobacterium bullatum* isolate mbul1 genome assembly
AGGCTGCCGGCTCGGTTGCCGGCTCAAGTCAGGATGGCTACATCGACCATCCCCGCAGCCGATGCGT
CCGATGGTTTCGCGAACCCTCTCGGACTGGCGTGGTTGCAGGGCTTGTCGTGACGGGGCGGTGCGGA
CCAGTTCTCTGCGCCGACCCTGTGAGGCCTCGTGGGCCGGGTCGCGGCAAGTGTGTCACGGGCGCGG
GGAACAGGAACCGTCGCCGTGGCTGATGGCCCACGCTCATCGTGCCAACGGACGGGCCGCGCCATTT
CAGGACACGCTCCAGGCCGGCCAGGGGCCAAATGGTTTACCGACTTCTCGCACAAGTGGGATGGGC
CGGGGAACCGGTCGCGTTCCTCCGCGTAGTCTCGGGCTTTCCAGCA
>*Methylobacterium bullatum* isolate mbul2 genome assembly
ATGCAGCCGGGTTGGTTGCCGGCTCAAGTCAGGATGTCTACATCCACCATCCCCGCAGCCGATGCGT
CCGATGGTTTCGCGAACCCTCTCGGACTGGCGCGGTTGCAGGGCTTGTCGTGACGGGGCGGTGCGGA
CCAGTTCTCTGCGTCGATCCTGTGAGGCCTCGTGGGCCGGGTCGCGGCAAGTGTGTCACGGGCGCGG
GGAACAGGAACCGTCGCCGTGGCTGATGGCCCACGCTCATCGTACTGGCGGACGGGTCGCGCCATTT
CAGGACAGGCTCCGAGCCGGCCAGCGGCCAAATGGTTTACCGACTTCTCGCACAAGTGAGATGGGC
CGGGGAACCGGTCGCGTTCCTCCGCGTAGTCTCGGGCTTTCCAGC
> Methylobacterium brachiatum strain TX0642
ATCGCCGCGGGTGGGCCGAATCGGCCGCCGCTCTGTTGCCGGCTCGGCTGCCACCCTCTACACGGAC

GGTCCCCGCGGCCGATGCGCCTGAAGGTTTCGCGAGCCCGCTCGGGCCTGCATGGTAGCGGGGCCGG
CAGTGACGGGGTGGTCCGGACCAGTTCTCTGGGCCAATCCTGTGAGGCTCAGTGGGCCGAGTCGCTG
CAGGAGTGTCGGGAGCGCGGGGAACAGGAACCGCGGGCTCGGCGCAAGGCCCACATAGAATTACCC
CGATCCGGGACTGGCGGCGCCGATCCGGCGCCCTGCCCCGATCCGGGACAGCCCGGTCGGGCGCGG
TTGCAGCTTCGTTCACGCTCCGGCGCGATGATACGACGAGGCCGGGGATCCGGTCCGGCGTAACCGC
GTTCTTTCGGGCCTGACGACCA
> *Methylobacterium mesophilicum* SR1.6/6
ATCGCCGCGGGGTGCGGGGAAACGGCTCCGCCTCGGTTGCCGGCTCGGCTGTCACCCTCTACACGGG
CTGTCCCCAGCGGCCGATGCGCCTGAAGGTTTCGCGAGCCCGCTCGGGCCTGCATGGTAGCGGGGTC
GGCAGTGACGGGGTGGTCCGGACCAGTTCTCTGGGCCGTTCCTGTGAGGCTCAGTGGGCCGAGTTGC
TGCAGGAGTGTCGGGCGCGCGGGGAACAGGAACCGCGGATCCGGCGCATGGCCCACATCAATCTCC
TTCCGAGATGGATCGATCCGCCCGCCCGGACCGTGCCCCGACGGGTCGGATGCCCCGTTCCGGGACA
GGGCCGGCGGCGGCGGTTGCGGCTTCGTTCACGCTCCCGCGGGATGATGCGGGAAGACCGGGGATC
CGGTCCTGCGTAACCGCGTTTTCTCGGGCCTGACGATCA
>Methylobacterium sp. C1
CCTCTACACGGGCTGTCCCCAGCGGCCGATGCGCCTGAAGGTTTCGCGAGCCCGCTCGGGCCTGCAT
GGCGGCGGGGTCGGCAGTGACGGGGCGGTCCGGACCAGTTCTCTGGACCGTTCCTGTGAGGCTCAGT
GGGCCGAGTCGCTGCAGGAGTGTCGGGAGCGCGGGGAACAGGAACCGCGGGTCCGGCGCATGGCCC
AC
>*Methylobacterium radiotolerans* JCM 2831
ATCGCCGCGGGGCCGCCGAAACGGCCCCGCCTCTGTTGCCGGCTCGGCTGTCACCCTCTACACGGGC
TGTCCCCAGCGGCCGATGCGCCTGAAGGTTTCGCGAGCCCGCTCGGGCCTGCATGGCGGCGGGGTCG
GCAGTGACGGGGCGGTCCGGACCAGTTCTCTGGACCGTTCCTGTGAGGCTCAGTGGGCCGAGTCGCT
GCAGGAGTGTCGGGAGCGCGGGGAACAGGAACCGCGGGTCCGGCGCATGGCCCACATGATTTCCCG
CCCGATCCCGCCCGCCGGACTTGTCGCGGGCGCGGGGCCCTGCCCCGTTCCGGGACAGGGGCGGCCG
GGGCCGTTGCGGCTTCGTTCACGCTCCCGCGCGATGATGCGTCCGGGCCGGGGATCCGGTTCCGCGT
AACCGCGTTCTCTCGGGCCTGACGACCA
>*Methylobacterium* sp. XJLW
GGGCCCTCCGGGCGATCCTTCCCGGGCGCGGACCCCGGTCCGCGCTCGGCGTGACGGGGCTCGCGCC
CGCGCTCTTGCCCGCGCTCTTGCCCGCGCTCTCGGGGGCGCCCTCCGGTCGGCCGCGGGGCCCGCTG
ATACGCTTCGCCGCGGCCGCGGCGGCGCGCGGATCGCCGCGCGGGCCACCGAAACGGTCCCGCCTCT
GTTGCCGGCTCGGCTGTCACCCTCTACACGGGCTGTCCCCAGCGGCCGATGCGCCTGAAGGTTTCGC
GAGCCCGCTCGGGCCTGCATGGCAGCGGGGTCGGCAGTGACGGGGCGGTCCGGACCAGTTCTCTGG
ACCGTTCCTGTGAGGCTCAGTGGGCCGAGTCGCTGCAGGAGTGTCGGGAGCGCGGGGAACAGGAAC
CGCGGGTTCGGCGCAAGGCCCACAAGATTCCCCGTCGACCGAGCGCCGGAGCCGTCCCGACGCCGC
GATCTGACCCGTTCCGGGACAGGGCCGAACGCGACGGTTGCGGCTTCGTTCACGCTCGCGCGCGATG
ATGCGTCCGGGCCGGGGATCCGGTTCCGCGTAACCGCGTTCTCTCGGGCCTGACGACCA
>*Methylobacterium phyllosphaerae* strain CBMB27
GGGCCCTCCGGGCGATCCTTCCCGGGCGCGGACCCCGGTCCGCGCTCGGCGTGACGGGGCTCGCGCC
CGCGCTCTTGCCCGCGCTCTCGGGGGCGCCCTCCGGTCGGCCGCGGGGCCCGCTGATACGCTTCGCC
GCGGCCGCGGCGGCGCGCGGATCGCCGCGCGGGCCACCGAAACGGTCCCGCCTCTGTTGCCGGCTC
GGCTGTCACCCTCTACACGGGCTGTCCCCAGCGGCCGATGCGCCTGAAGGTTTCGCGAGCCCGCTCG
GGCCTGCATGGCAGCGGGGTCGGCAGTGACGGGGCGGTCCGGACCAGTTCTCTGGACCGTTCCTGTG
AGGCTCAGTGGGCCGAGTCGCTGCAGGAGTGTCGGGAGCGCGGGGAACAGGAACCGCGGGTTCGGC
GCAAGGCCCACAAGATTCCCCGTCGACCGAGCGCCGGAGCCGTCCCGACGCCGCGATCTGACCCGTT
CCGGGACAGGCCCGAACGCGACGGTTGCGGCTTCGTTCACGCTCCCGCGCGATGATGCGTCCGGGCC
GGGGATCCGGTTCCGCGTAACCGCGTTCTCTCGGGCCTGACGACCA
>Methylobacterium oryzae CBMB20
GGGCCCTCCGGGCGATCCTTCCCGGGCGCGGACCCCGGTCCGCGCTCGGCGTGACGGGGCTCGCGCC
CGCGCTCTTGCCCGCGCTCTTGCCCGCGCTCTTGCCCGCGCTCTCGGGGGCGCCCTCCGGTCGGCCGC
GGGGCCCGCTGATACGCTTCGCCGCGGCCGCGGCGGCGCGCGGATCGCCGCGCGGGCCACCGAAAC
GGTCCCGCCTCTGTTGCCGGCTCGGCTGTCACCCTCTACACGGGCTGTCCCCAGCGGCCGATGCGCCT
GAAGGTTTCGCGAGCCCGCTCGGGCCTGCATGGCAGCGGGGTCGGCAGTGACGGGGCGGTCCGGAC
CAGTTCTCTGGACCGTTCCTGTGAGGCTCAGTGGGCCGAGTCGCTGCAGGAGTGTCGGGAGCGCGGG
GAACAGGAACCGCGGGTTCGGCGCAAGGCCCACAAGATTCCCCGTCGACCGAGCGCCGGAGCCGTC
CCGACGCCGCGATCTGACCCGTTCCGGGACAGGGCCGAACGCGACGGTTGCGGCTTCGTTCACGCTC
GCGCGCGATGATGCGTCCGGGCCGGGGATCCGGTTCCGCGTAACCGCGTTCTCTCGGGCCTGACGAC
CA
>*Methylobacterium radiodurans* strain 17Sr1-43
GGCCACGAGTTGTGACGCAGAGAGCATCACGGTCGAACCGCCGAGCTTGTTTCCCCCGGTTATCGGG
GCCTGAGATGGAGCCAGGACGGATCGCCGCGGTTGCTTGGCGCCCGCGCGCCCTCTACATCGGCGGT
CCCCGCGGCCGATGCGCCTGAAGGTTTCGCGAGCCCGCTCGGGCCGGCATGGTAGCGGGGTCGGCA
GCGACGGGGCGGTCCGGACCAGTTCTCTGGGCCGCTCCCGTGAGACCCAGTGGGCCGGGCGGCTGC
AGGGGTGTCGGAGGCGCGGGGAACAGGAACCGCGGATCCGGCGCATGGCCCACAAACTTCTCCGGG
CTCGTCCATGATCTGGCGCGTGATCTGACCCGCGCCTGAGCTGGCCCGTTCCGGGACGAAGACACCG
GCAGGTTCTTGTGGCTTCGTTCACGCCGCCCGGTAAGGATAAGGACGGGCCGGGGAACCGGCCGCGT
GCCACCGCGTTCCTTCGGGCCTGACGACCAT

Supplementary material

>*Methylobacterium* sp. WL1
ATCGCCGCGGGGGCGCCGAATCGGCCGTCGCTCTGTTGCCGGCTTGGCTGCCATTCTCTACACGAGC
GGTCTCCCGCGGCCGATGCGCCTGAAGGTTTCGCGAGCCCGCTCGGGCCTGCATGGTAGCGGGGCCG
GCAGTGACGGGGTGGTTCGGACCAGTTCTCTGGACCAATCCTGTGAGGCTCAGTGGGCCGAGTCGCT
GCAGGAGTGTCGGGAGCGCGGGGAACAGGAACCGCGGGCTCGGCGCAAGGCCCACAAAGATTTCCG
TTCGCTATCGGCGTCGACAGGTGCCCGCGGCGGCATCCGGGCAGCCCGATCCGACGCCTCGCCCGAT
CCGGCGCCCTGCCCCGATCCGGGACGATGCAACGCCAGGCGGTTGCAGCTTCGTTCACGCTCCCGCG
CGATGATGCGATAGGGCCGGGATCCGGTCCGGCGTAACCGCGTTCTTTCGGGCCTGACCACCA
>*Methylobacterium nodulans* ORS 2060
AAGTAGCCCCGCCGCGTTTCGACGCAGCCGAGCGCGACCATTCCTGTTGCCCCGCCGGCGTGGTCTG
CTACACGGGCAGTCCTGACCGCCAGCGCGCCCGGAGGCTTCCGGAACCCGTCTGGGCTGCGCGGTGG
AAGGGCCGGCGATGACGGGATGGGGCGGACCAGTTCTCTGCAACCCATTCCTGTGAGGCTCAGTGG
GCCGAGTAGTCGCAGGAGCGTCAGTGGTTCGGGGAACAGGAACCGCGACCCTGGCGCATGGCCCAC
CAGGATTCACCCGCATGAGCCGGCGCGGCGATCGGGAGACGCCGGCATCCGCCTTTGCGAAGCAAT
CTGCCGGATCCTGTATGGCACCCTTGTTAACGGGTGCGGCCTAGGGTCGGCGGGTCGAATTCCCCAG
GCGAGTGTCCGGCCA
>*Methylobacterium currus* strain PR1016A
ACCCGGCCTCACGGCCGAGTGCCTGATTCGACCCGGCGGGCCCGTGCCCGCCGGGTTTTTCGTGAGC
GCGGTCAAGCTTTCCGGCCGGCGCGTTTAGACGCAAGCGCGGAGGCCCCATTCCGGTTGCCCCGCGG
CCCCGGTCTGGCTTACACGGGCCGTCCTCGATGCCGGCGCGCCCGGAGGCTTCCGGAACCCATCCGG
GCTGTGCGGCAAAGAGGTCGGCGATGACGGGGTGGGGCGGACCAGTTCTCTGCAACCCATTCCTGTG
AGGCTCAGTGGGCCGAGCAGTCGCAGGAGCGTCACGGGTTCGGGGAACAGGAACCGCGACCCTGGC
GCATGGCCCACCAGAGTTTCCGACGACTCGTCGGACTCCGCTCGTCGGCTCCCGTCGTCTTGGCGGC
CCGAGGCTCCCGTTGTGGGACGTCCTGAGATCAGCTGCGAGGGGCCGCGCGTTAAGCATTGTGGCCC
AAGACTAGCGGGACGGGCCGGATCGGCGGCCCGGCGAGATCGTGGGTCCCGACCAGCCA
>*Methylobacterium* sp. 17Sr1-1
ACGCGCAGATCCAGAGGCCTCGCGTGATTCGACCCGGCGGGCCCGTGCCCGCCGGGTTTTTCGTTGC
GCCGGTCAAGCTTTCCCGGCCGACGCGTTTAGACGCAAGCGCCGCCCCTGCATTCCGGTTGCGACCC
CGACCCGGTCTGGCTTACACGGACCGTCCTCGATGCCGGCGCGCCCGGAGGCTTCCGGAACCCATCC
GGGCTGTGCGGCAAAGAGGTCGGCGATGACGGGGTGGGGCGGACCAGTTCTCTGCAACCATTCCTG
TGAGGCTCAGTGGGCCGAGCAGTCGCAGGAGCGTCACGGGTTCGGGGAACAGGAACCGCGACCCTG
GCGCATGGCCCACCAAACTTCGCCTCTTTCGCCGATCGGCCGCTCCCGTTGCGGGACAGTCCCGGCG
AGCGGGCCGAGGCCGCCCGTTAAGCATTGTGGCCCAGGATCGGCGAAACGGGGCCGGATCGACGGC
CCCGACGAAATCGCGGGTCCCGACCAGCCA
>*Methylobacterium indicum* VL1 DNA
ATCGCTCCGCGAGCCGGCCGCGCCTGCGCGGCCGGATGACGAGGTGTTCGAACCCGGCGGGCCCTG
GGCCCGCCGGGTTTATCGTCTCGGTCGGTCAAGTGCGCCGGGCCGGCGCGTTGTGACGCAAGCGCCT
CGGCGGTATTCCGGTTGCCCCTCCGGCCCCGTCTGTCCTACACGGGCCGTCCTCGATGCCGGCGCGCC
CGGCGGCTTCCGGAACCCGTCCGGGCTGTGCGGCAAAGAGGTCGGCGACGACGGGGTGGGGCGGAC
CAGTTCTCTGCACCCAATCCTGTGAGGCTCAGTGGGCCGAGCAGTCGCCGGAGCGTCAGGGTTCCGG
GGAACAGGAACCGCGATCCTGGCGCAAGGCCCACCAGATTGCTCCGCCCTCCCGTTGTGGGACGATC
CGGGTCGGGGGCGGGGAGGCCGCCCGTTAAGGCTTATGGCCCAAGAATGTCGGACGGCCGGATCGT
CGGCCTCACACCGATCGTGGGTCCCGACCAGCCA
>*Methylobacterium aquaticum* strain BG2
AACAGGCCCTAGGCCGATCGCGTGATTCGACCCGGCGGGCCCTGAGCCCGCCGGTTTTTCTTGGGG
CCCGTCAAGCTTTTCCGGCCGGCGCGTTTCGACGCAAGGGCCAGGGCCGCATTCCGGTTGCCCCACC
CGCCCGGTCTGGCTTACACGGGCCGTCCTCGATGCCGGCGCGTCCGGAGGCTTCCGGAACCCATCCG
GGCTGTGCGGCAAAGAGGTCGGCGACGACGGGATGGGGCGGACCAGTTCTCTGCACCCATTCCTGT
GAGGCTCAGTGGGCCGAGCAGTCGCAGGAGCGTCACGGGTTCGGGGAACAGGAACCGCGACCCTGG
CGCATGGCCCACCAGACTTTTCGGGCATCGCTCCGGGCAATCCGGAATATTGAGCGCCGTCGATCCG
TCATCCCGTGTTCCGCTGTCGCGGCGCCGGGATGACGGGGGAGAATTCGAGCGCGCCCGACCCGATC
GAGCGGCCCCCGGATCGACCGCCTGCTGCCCCTCGACCTCGTTCGACCCGTCTTCATCGGAACTCCC
GTTGTGGGACGACCCGATCCGCGCCGCGAGGGGCCACGCGTTAAGCATTGTGGTCCGATACTGTCGG
CATGGGCCGGACCGACGGCCCACCGAGATCACGGGTCCCGACCAGTCA
> *Methylobacterium aquaticum* strain MA-22A
AGCGCGCCGCGCTGCAGGATGGCGGGATGAATCGACCCGGCGGGCCAAGTCCCCGCCGGGTTTTCT
GCCCCTGGGCTCTTGGCCTCCTCCGGTCAAGCGCACGTGGCCGGCGCGTTTCGACGCAAGGCCGCCG
CCCGCATTCCGGTTGCCCCTCCGGCCCGGTCTGTCCTACACGGACCGTCCTCGATGCCGGCGCGCCCG
GTGGCTTCCGGAACCCGTCCGGGCTGTGCGGCAAAGAGGTCGGCGATGACGGGGTGGGGCGGACCA
GTTCTCTGCAACTCATTCCTGTGAGGCTCAGTGGGCCGAGCAGTCGCCGGAGCGTCAGGGTTCCGGG
GAACAGGAACCGCGATCCTGGCGCAAGGCCCACCAGACTTGTCCCGCCGGCCTCGCCCGCTCTCCCG
ATGTGGGACGGCCCGCCCCGGGACCGTGAAGGCCGCCCGTTAAGTCCTGTGGCTCAAGGATGTCGG
GACGGGCCGGATTGCCGGCCGCACACCGATCGTGGGTCTCGACCAGCCA

Supplementary material

| Hits from Metagenomic data |
|---|

>PJQF01118147.1
GGTAAGCGGCGCTCCGATCGTTTTGGCAATGCATTGCCGGTAGGGCAGCGGCGTCAGGGCGCGGTTG
CCCGCGGCGGCGCGGGCCTCTACATCGGCCCTCCCCGCGGCCGATGCGCCCGATGGTTTCGCGAGCC
CTCTCGGGCCTGCGTGGCGGCGGGGCCGGCAGTGACGGGGCGGTGCGGACCAGTTCTCTGCGCCGTT
CCTGTGACGCTCAGTGGGCCGAGCGGCTGCCGGTGCGTCGGGAGCGCGGGGAACAGGAACCGCGAT
CCCGGCGCAAGGCCCACCTATTCGGCTCCCGTCAGGCCGAAGAGGTCACCCGGCCCTGTGCCGGAGT
GCCGTTAACGGTCTTTCTCAAGGGAAAGCGAACCACTCAGAT

> FUFK012632339.1
ACGCCACCTCCAACAAGACCCTGGAGGAGGCCTGCCGGCGCATCCAGCGCTTCTGCGGCTCGCTGCG
GTAAGCGGCGCTCCGATCGTTTTGGCAATGCATTGCCGGTAGGGCAGCGGCGTCAGGGCGCGGTTGC
CCGCGGCGGCGCGGGCCTCTACATCGGCCCTCCCCGCGGCCGATGCGCCCGATGGTTTCGCGAGCCC
TCTCGGGCCTGCGTGGCGGCGGGGCCGGCAGTGACGGGGCGGTGCGGACCAGTTCTCTGCGCCGTTC
CTGTGACGCTCAGTGGGCCGAGCGGCTGCCGGTGCGTCGGGAGCGCGGGGAACAGGAACCGCGATC
CCGGCGCAAGGCCCACCTATTCGGCTCCCGTCAGGCCGAAGAGGTCACCCGGCCCTGTGCCGGAGTG
CCGTTAACGGTCTTTCTCAAGGGAAAGC

> FUFK010110014.1
ATGGGGCGCGGCTCCCGGCGGGTTTGAATGCGGCGCAGCATGGGTTACACCCGGCGCCGTTCAGAA
GACCTCATCCGGGAGCCAAATCCTCATGTCCCTGGAATCCGCCGCGCTCGCCCGCGTGAAGCCGTCC
GCGACCCTCGCGGCCGACGCGAAGGCCCGCGAGCTGAAAGCCCAGGGCAAGGACGTCATCGGCCTG
GCCGCCGGCGAGCCCGACTTCGACACGCCGGACAACATCAAGGACGCCGCCATCAAGGCGATCCGC
GACGGCAAGACCAAGTACACCAACGTCGATGGCATCCCGGAGCTGAAGGAGGCGATCTGCGCCAAG
TTCCACCGGGAGAACGGCCTTTCCTACAAGCCCAGCCAGATCAACGTCTCGCCGGGCGGCAAGCCGG
TGATCTGGAACGCCATGATCGCCACGCTGAACCCCGGCGACGAGGTGATCGTCCCCACGCCCTACTG
GGTCAGCTACTGGGACATCGTGCTGCTGGCCGGGGGGACGCCCGTCGCCCTGCCCAGCTCAGCCGAC
GTCGGCTTCAAGGTCCAGCCGGCGGATCTGGAGCGTGCGATCACGCCGAAGACCAAGTGGATCATC
CTCAACTCGCCGTCGAACCCGTCGGGCGCGGCCTATACGCGCGCCGAGCTGCGCGCCCTGGCCGACG
TGCTGCTTAAGCACCCGCATGTCTGGATCCTGACCGACGACATGTACGAACACCTGGTGTTCGGCGA
CTTCGAGTTCACCACCATCGCCCAGGTCGAGCCCGGCCTCTACGACCGCACCCTGACCATGAACGGG
GTCTCNAAGGCCTACGCCATGACCGGCTGGCGCATCGGCTACGCCGCGGGCCCCGAACAGCTCATCA
AGGCGATGGACTTCGTGCAGGGCCAGCAGACCTCGGGCGCCTCCTCGATCTCGCAATGGGCGGCGGT
GGCGGCGCTCGACGGGACGCAGGAGCACCTCGCCCGGTTCAAGGCCGCGTTCCAGGAGCGGCGCGA
CCTCGTGGTCTCGATGCTCAACCAGTCGAACGGCCTGAAATGCCCGGTGCCGGAGGGCGCGTTCTAC
GTCTATCCGTCCTGCGCCGATCTGATCGGCAAGACCACCGAGACCGGCAAGACCATCGCCACGGACG
AGGATTTCGTCACCGAGCTGCTCCAGGCGGAGGGTGTCGCCGCGGTGCACGGCTCGGCCTTCGGCCT
CGGCCCGAACCTGCGCATCTCCTACGCCACCTCCAACAAGACCCTGGAGGAGGCCTGCCGGCGCATC
CAGCGCTTCTGCGGCTCGCTGCGGTAAGCGGCGCTCCGATCGTTTTGGCAATGCATTGCCGGTAGGG
CAGCGGCGTCAGGGCGCGGTTGCCNGCGGCGNCGCGGNCCTCTACATCGGCCCTCCCCGCGGCCGAT
GCGCCCGATGGTTTCGCGAGCCCTCTCGGGCCTGCGTGGCGGCGGGGCCGGCAGTGACGGGGCGGT
GCGGACCAGTTCTCTGCGCCGTTCCTGTGACGCTCAGTGGGCCGAGCGGCTGCCGGTGCGTCGGGAG
CGCGGGGAACAGGAACCGCGATCCCGGCGCAAGGCCCACCTATTCGGCTCCCGTCAGGCCGAAGAG
GTCACCCGGCCCTGTGCCGGAGTGCCGTTAACGGTCTTTCTCAAGGGAAAGCGAACCACTCGCGGGC
AATATTTCGCCAGTTGTCCAACAGAGGTCGCTCCATGTCTCACGCCGGCAGAGCCCGCGCTTCCGCC
CGCCAGTACCTGCCGGAATCGAAGCTCGAGGATCTCGCCAGCAGCCTGCGCCGGCTCACCAATCATC
GCGGCCTCGTGCGCAG

> FUFK010045813.1
ACGGACGAGGATTTCGTCACCGAGCTGCTCCAGGCGGAGGGTGTCGCCGCGGTGCACGGCTCGGCCT
TCGGCCTCGGCCCGAACCTGCGCATCTCCTACGCCACCTCCAACAAGACCCTGGAGGAGGCCTGCCG
GCGCATCCAGCGCTTCTGCGGCTCGCTGCGGTAAGCGGCGCTCCGATCGTTTTGGCAATGCATTGCC
GGTAGGGCAGCGGCGTCAGGGCGCGGTTGCCCGCGGCGACGCGGACCTCTACATCGGCCCTCCCCG
CGGCCGATGCGCCCGATGGTTTCGCGAGCCCTCTCGGGCCTGCGTGGCGGCGGGGCCGGCAGTGACG
GGGCGGTGCGGACCAGTTCTCTGCGCCGTTCCTGTGACGCTCAGTGGGCCGAGCGGCTGCCGGTGCG
TCGGGAGCGCGGGGAACAGGAACCGCGATCCCGGCGCAAGGCCCACCTATTCGGCTCCCGTCAGGC
CGAAGAGGTCACCCGGCCCTGTGCCGGAGTGCCGTTAACGGTCTTTCTCAAGGGAAAGCGAACCACT
CGCGGGCAATATTTCGCCAGTTGTCCAACAGAGGTCGCTCCATGTCTCACGCCGGCAGAGCCCGCGC
TTCCGCCCGCCAGTACCTGCCGGAATCGAAGCTCGAGGATCTCGCCAGCAGCCTGCGCCGGCTCGCC
AATCATCGCGGCCTCGTGCGCAGCGAGATCGCCAGCCCGATGCTGCTGCGTCTGCTGCCGCCCCCGC
GGCGCGCCGAGCAGAAGAGCTACGAGGCCGACCTGCGCCAGCGCCTGACCGACGCCAACCTCGATG
GGTCCCGCATCGCCTACCTGATGGCCGATGCCGAGCGGGAGATCGCGATCGCCCATACCCGCCTGTC
GGGCTGAGCCGGCCGCGGTGGTCGGGAACCCACCGCGACATGAACCAAGGGTTGGAGCGCGGCTC

#### Table S3. Intergenic regions containing Met1153.

More conserved region containing Met1153 based on alignment results of all intergenic regions are italicized.

**Hits from Genomic data**

> Methylobacterium extorquens AM1
ATCGCCTGTTCGGCGAATGGGGTCAGCCGGTTGGTGGATCAGATTCGCTTCTCCGGCCAATCCCCTCA
TCCTGAGGTGCCGGAGCGAAGCGGAGGCCTCGAAGGAGGGCTCCAGAAGACGCTGCGATTCCTGGA
GCCCTCCTTCGAGGCCGCTCCGCGGCACCTCAGGATGAGGTAAGTCCTGGGAAAACCAGTCGAATAG
AATCTCAAGCGAACGGCACGATCAGCGGCAGCGCGAGCAGGGCGCCGGCGGCCAGGACGATGGCAT
CGCGCAGGCGCCGGCCGATCCAGGCGCGGGCGTCGGGATGCTGTGCGTGGGTGTCTTCGAAAAACGT
CATGGTCTGTCTCCTGTCGGTTCGATGGCCCCGATATTACGGCGCCGCCGCTTTCCCGACAGGGACGG
TTGCGGTACGATTTCAACGAACTGTCCGGGACGAGCACCGATACAGTTCGGAGGGACAGCCA
>Methylorubrum zatmanii strain PSBB041
CATGGAGCGGGCGAAGGCGCTGTTCGAGGCGAACTGCACCGCCGGCTGATCGCCTGTTCGGCGAATG
GGGTCAGCCGGTTGGTGGATCAGATTCGCTCCTCCGGCCAATCCCCTCATCCTGAGGTGCCGGAGCGA
AGCGGAGGCCTGGAAGGAGGGCTCCAGAAGACGCTGCGATTCCTGGAGCCCTCCTTCGAGGCCGCTC
CGCGGCACCTCAGGATGAGGTAAGTCCTGGGAAAACCAGTCGAATAGAATCTCAGGCGAACGGCAC
GATCAGCGGCAGCGCGAGCAGGGCGCCGGCGGCGAGGACGATGGCATCGCGCAGGCGCCGGCCGAT
CCAGGCGCGGGCATCGGGATGCTGTGCGTGGGTGTCTTCAAAAAACGTCATGGTCTGTCTCCTGTCGG
TTCGATGGCCCCGATATGACGGCGCCGCCGCTTTCCCGACAGGGACGGTTGCGGTACGATTTCAACGA
ACTGTCCGGGACGAGCACCGGTACAGTTCGGAGGGACAGCCATGACGGTGACGGTCGCGGAACCGGT
CCTCTCGCGGGTCGAGACGGTGATG
>Methylobacterium extorquens strain TK 0001
GAGCGAAGCGGAGGCCTGGAAGGAGGGCTCCAGAAGACGCTGCGATTCCTGGAGCCCTCCTTCGAGG
CCGCTCCGCGGCACCTCAGGATGAGGTAAGTCCTGGGAAAACCAGTCGAATAGAATCTCAGGCGAAC
GGCACGATCAGCGGCAGCGCGAGCAGGGCGCCGGCGGCGAGGACGATGGCATCGCGCAGGCGCCGG
CCGATCCAGGCGCGGGCATCGGGATGCTGTGCGTGGGTGTCTTCAAAAAACGTCATGATCTGTCTCCT
GTCGGTTCGATGGCCCCGATATGACGGCGCCGCCGCTTTCCCGACAGGGACGGTTGCGGTACGATTTC
AACGAACTGTCCGGGACGAGCACCGGTACAGTTCGGAGTGACAGCCATGACGGTGACGGTCGCGGA
ACCGGTCCTCTCGCGGGTCGAGACGGTGATGCGGGCCATCGAGGCGCGGATCGAGGGCCGGGCGCTC
GGCCCCGGCGCGCGGCTGCCCTCGGTGCGGAGCCTCGCCGACA
>METHYLORUBRUM EXTORQUENS STRAIN PSBB040
TGGCTGCCCCTCCGAACTGTACCGATGCTCGTCCCGGACAGTTCGTTGAAATCGTACCGCAACCGTCC
CTGTCGGGAAAGCGGCGGCGCCGTCATATCGGGGCCATCGAACCGACAGGAGACAGACCATGACGTT
TTTCGAAGACACCCACGCCCAGCATCCCGACGCCCGCGCCTGGATCGGTCGGCGCCTGCGCGATGCC
ATCGTCCTCGCCGCCGGCGCCCTGCTCGCGCTGCCGCTGATCGTGCCGTTCGCCTGAGATTCTATTCG
ACTGGTTTTCCCAGGACTTACCTCATCCTGAGATGCCGCGGAGCGGCCTCGAAGGAGGGCTCCAGGA
ATCGCAGCGTCTTCTGGAGCCCTCCTTCGAGGCCTCCGCTTCGCTCCGGCACCTCAGGATGAGGGGAT
TGGCCGGAGGAGCGAATCTGATCCACCAACCGGCTGACCCCATTCGCCGAACACGCGATCAGCCGGC
GGTGCAGTTCGCCTCGAACAGCGCCTTCGCCCGCTCCATGGCCCGGAACGCCGCCACGGAGCCCGGG
CTGCGCAGTTCCGGCCCCTCGGGCAGACGCCCCAGGAAGGGTTCGGACAGGTCCGCCACGAACGGCC
GGGGCCCGGTATAGGCGCCCATGCGGTTGCGCAGATCGACCGTGCCGCACAAGGCCCCGGCCCGGCC
GATCCGCAGGCCGGCCACCTTCGCCTCGGGGTCGCGCATCTGCCGCTCGATCAGGCTCAGCGCGGTCC
GCGCCGCCTCCGCCCCGACGCCGGCCTCCGAACCGTCCACGACGGTCTGAGCGCCCGCCGCACCCGC
GCCGGGCAGGATCAGGAAGGCCGCGAGGGCGGTCTTGCGCAGGCGCAT
>Methylobacterium extorquens CM4
GAGGCGAACTGCACCGCCGGCTGATCGCCTGTTCGGCGAATGGGGTCAGCCGGTTGGTGGATCAGAT
TCGCTCCTCCGGCCAATCCCCTCATCCTGAGGTGCCGGAGCGAAGCGGAGGCCTCGAAGGAGGGCTC
CAGAAGACGCTGCGATTCCTGGAGCCCTCCTTCGAGGCCGCTCCGCGGCACCTCAGGATGAGGTAAG
TCCTGGGAAAACCAGTCGAATAGAATCTCAGGCGAACGGCACGATCAGCGGTAGCGCGAGCAGGGC
GCCGGCGGCCAGGACGATGGCATCGCGCAGGCGCCGGCCGATCCAGGCGCGGGCACCAGGATGCTG
TGCTTGGGTGTCTTCAAAAAACGTCATGGTCTGTCTCCTGTCGGTTCGATGGCCCCGATATGACGGCG
CCGCCGCTTTCCCGACAGGGACGGTTGCGGTACGATTTCAACGAACTGTCCGGGACGAGCGCTGGTA
CAGTTCGGAGAGACAGTCATGACGGTGACAGTCGCGGAACCGGT
>Methylobacterium extorquens str. DM4 chromosome
GAGGCGAACTGCACCGCCGGCTGATCGCGTGTTCGGCGAATAGGGTCAGCCGGTTGGTGGATCAGAT
TCGCTCCTCCGGCCAATCCCCTCATCCTGAGGTGCCGGAGCGAAGCGGAGACCTCGAAGGAGGGCTC
CAGAAGACGCTGCGATCTCTGGAGCCCTCCTTCGAGGCCGCTCCGCGGCACCTTAGGATGAGGTAAG
TCCTGGGGAAAACCCAGTCGAAGAGCCGCTCAGGCGAACGGCACGATCAGCGGCAGCGCGAGCAGG
GCGCCAGCGGCGAGGACGATGGCATCGCGCAGGCGCCGGCCGATCCAGGCGCGGGCGTCGGGATGC
TGTGCGTGGGTGTCTTCGAAAAACGTCATGGTCTGTCTCCTGTCGGTTCGATGGCCCCGATATGACGG
CGCCGCCGCTTTCGCGACAGGGACGGTTGCGGTACGATTTCAACGAACTGTCCGGGACAAGCGTCGG
TACAGTTCGGAGGGACATCCATGACGGTGACAGTCGCGGAACCGGT

### Supplementary material

>Methylobacterium sp. AMS5
ATCGCGCAGAGCCGAATCCGGCCCGATCGCGTCAGCCCGGCGTCTCAAGGTCCTTGCTTTCGCACCGT
CGCTTTCCGAAAGCCGGCGACCGCCTTTCGGGGCGATGCCCTAGGCGAACGGCACGATCAGCGGCAG
GGCGAGCACGGCGCCGGCGGCCAGGACGATGGCATCGCGCAGGCGCCGGCCGATCCAGGCGCGGGC
GCCGGGATGCGGCGCGTGGGTGTCTTCAAAAAACGTCATGGTCTGTCTCCTATCGGGCTCGATGGCAC
CGATATGACGGCGCCGCCGCTTTCCCGACAGGGACGGTTGCGGTACGATTTCGAAGAACTGTCCGGG
ACGAGCGCCGGTACAGTTCGAAGGGACAGCCA

>Methylobacterium populi BJ001
ACGGCCTCCTTGGCGACTCGGCTCAGCCAGCGAGTGGCTCAGATCCACTCGTTCAGCCAATCCCTCAC
CTGAGGTGACGGAGCGAAGCGGAGCCCTCCTTCGAGGCCTCTCCGCGGCACCTCCGGAAATATGAGG
TCCTGGGACAAGCCGGCTGATCGTCCCTCAAGCGAACGGCACGATCAGCGGCAGGGCCAGCGCGGCG
CCGGCGGCCAGGACGATGGCATCGCGCAGGCGGCGGCCGATCCAGGCGCGGGCGCCGGGCCGGCAT
GCAGGGGCGTCTTCTGCTTGGGTGTCTTCTGAAAGCGTCATGATCCGTCTCCCGTCTCTCGGTGCTCCC
GATATGACGGCGGCACCGCTTTCCCGACAGGGACGGTTGCGGTACGATTTCAAGGAACTGTCCGGGA
CATGGTTCGGTACAGTTCGGAGGGACAGTCA
>Methylorubrum populi strain YC-XJ1
ACGGCCTGCTTGGCGACTCGACTCAGCCAGCGCGTGGCTCAGATCCACTCGTTCAGCCAATCCCTCAC
CTGAGGTGACGGAGCGAAGCGGAGCCCTCCGTCGAGGCCGCTCCGCGGCACCTCAGGAACCATGAGG
TCCTGGCACAAGCCGGCTGATCGTCCCTCAAGCGAACGGCACGATCAGCGGCAGGGCCAGCGCGGCG
CCGGCGGCCAGGACGATGGCGTCGCGCAGGCGCCGGCCGATCCAGGCGCGGGTGCCGGGCCGGCAT
GCAGGGGCGTCTTCTGCTTGGGTGTCTTCTGAAAGCGTCATGATCCGTCTCCCGTCTCTCGGTGCTCCC
GATATGACGGCGGCACCGCTTCCCCGACAGGGACGGTTGCGGTACGATTTCAAGGAACTGTCCGGGA
CATGGTTCGGTACAGTTCGGAGAGACAGTCA
>Methylobacterium sp. NI91
AGTCGGCTCAGGCGAACGGCACGATCAGCGGCAGCGCCAGCACGGCGCCGAGGGCCAGGACGAGGA
CGTCCCGCAGGCGCCGGCCGAGCCAGGCTCGGGTGCCGGGTTGGCGCGAATGGGTGTCTTCTGAAAG
CGTCATGGTCTGTCTCCCGTCTTTCGGTGGCTCCGATATGAGACGATTGCCGCTTCTCCGACAGGGAC
GGTTGCGGTACGATTCCGACGAACTGTCCAGGACGAACGCCGGTACAGTTCGGAGGGACAATCA
>Methylobacterium sp. CLZ
AGTCGGCTCAGGCGAACGGCACGATCAGCGGCAGCGCCAGCACGGCGCCGAGGGCCAGGACGAGGA
CGTCCCGCAGGCGCCGGCCGAGCCAGGCTCGGGTGCCGGGTTGGCGCGAATGGGTGTCTTCTGAAAG
CGTCATGGTCTGTCTCCCGTCTTTCGGTGGCTCCGATATGAGACGATTGCCGCTTCTCCGACAGGGAC
GGTTGCGGTACGATTCCGACGAACTGTCCAGGACGAACGCCGGTACAGTTCGGAGGGACAATCA
>Methylobacterium sp. DM1
AGTCGGCTCAGGCGAACGGCACGATCAGCGGCAGGGCTAGCACGGCGCCGAGGGCCAGAACGAGGA
CGTCCCGCAGGCGCCGGCCGAGCCAGGCTCGGGTGCCGGGTTGGCGCGAATGGGTGTCTTCTGAAAG
CGTCATGGTCTGTCTCCCGTCTCTCGGTGGCTCCGATATGAGACGATTGCCGCTTCTCCGACAGGGAC
GGTTGCGGTACGATTTTGACGAACTGTCCAGGACGAGCGCCGGTACAGTTCGGAGGGACAATCA
>Methylorubrum sp. B1-46
AGTCGGCTCAGGCGAACGGCACGATCAGCGGCAGGGCCAGGAGAGCGCCGAGGGTCAAAACGAGGA
GGTCGCGCAGGCGCCGGCCGAGCCAGGCACCGGTGCCGGGCTGGCGCGAATGGGTGTCTTCTGAAAG
CGTCATGGTCAGTCTCCCGTCTCTCGGTGGCTCCGATATGAGGGGATCGCCGCTTCTCCGACAGGGAC
GGTTGCAGTACGATTCCGAGGAACTGTCCGGGGCATGCGCCGGTACAGTTCGGAGGAACAGTCA
>Methylobacterium populi DNA
AGCGGCCTCAGGCGAACGGCACGATCAGCGGCAGGGCGAGCGCGGCCCCGGCCGCCAGGACGATGG
CGTCGCGCAAGCGCCGGCCGATCCAGGCACGGGCGCCGGGACGGCGCGCGCGGGTGTCTTCTGAAAG
CGTCATGGTCTGTCTCCCATCTGACGGTCGCCCCGATATGCCGGCGGCGCCGCTTCCCCGACAGGGAC
AGTTGCAGTACGATTTCATCGAACTGTCCGGGACAAGCGTCGGTACAGTTCGGAGGAACAGTCA

**Table S4. RNAcode results for intergenic region containing Met2624.**

| Frame | Amino acids (aa) | | | Nucleotide | | Score | P |
|---|---|---|---|---|---|---|---|
| | Length | From | to | Start | End | | |
| -2 | 29 | 3 | 31 | 8 | 94 | 17.62 | 0.305 |
| +2 | 20 | 1 | 20 | 2 | 61 | 16.03 | 0.439 |
| -3 | 9 | 54 | 62 | 162 | 188 | 11.46 | 0.890 |
| +3 | 16 | 34 | 49 | 102 | 149 | 10.18 | 0.959 |
| -1 | 9 | 34 | 42 | 100 | 126 | 10.05 | 0.964 |
| +1 | 8 | 21 | 28 | 61 | 84 | 7.94 | 0.998 |
| +2 | 4 | 23 | 26 | 69 | 80 | 7.14 | 1 |
| -2 | 14 | 39 | 52 | 116 | 157 | 7.10 | 1 |
| +3 | 3 | 5 | 7 | 15 | 23 | 6.45 | 1 |
| +2 | 14 | 47 | 60 | 140 | 181 | 6.21 | 1 |
| -1 | 3 | 5 | 7 | 15 | 23 | 5.71 | 1 |

**Table S5*: Optical density at which samples were taken for RNA-seq***

| Condition | | Optical density (600 nm) |
|---|---|---|
| | 1 | 3.206 |
| Wild type, controlled pH | 2 | 3.12 |
| | 3 | 3.125 |
| | 1 | 2.968 |
| Triple mutant, controlled pH | 2 | 2.902 |
| | 3 | 2.921 |
| | 1 | 2.958 |
| Triple mutant | 2 | 3.266 |
| | 3 | 3.207 |

#### Figures

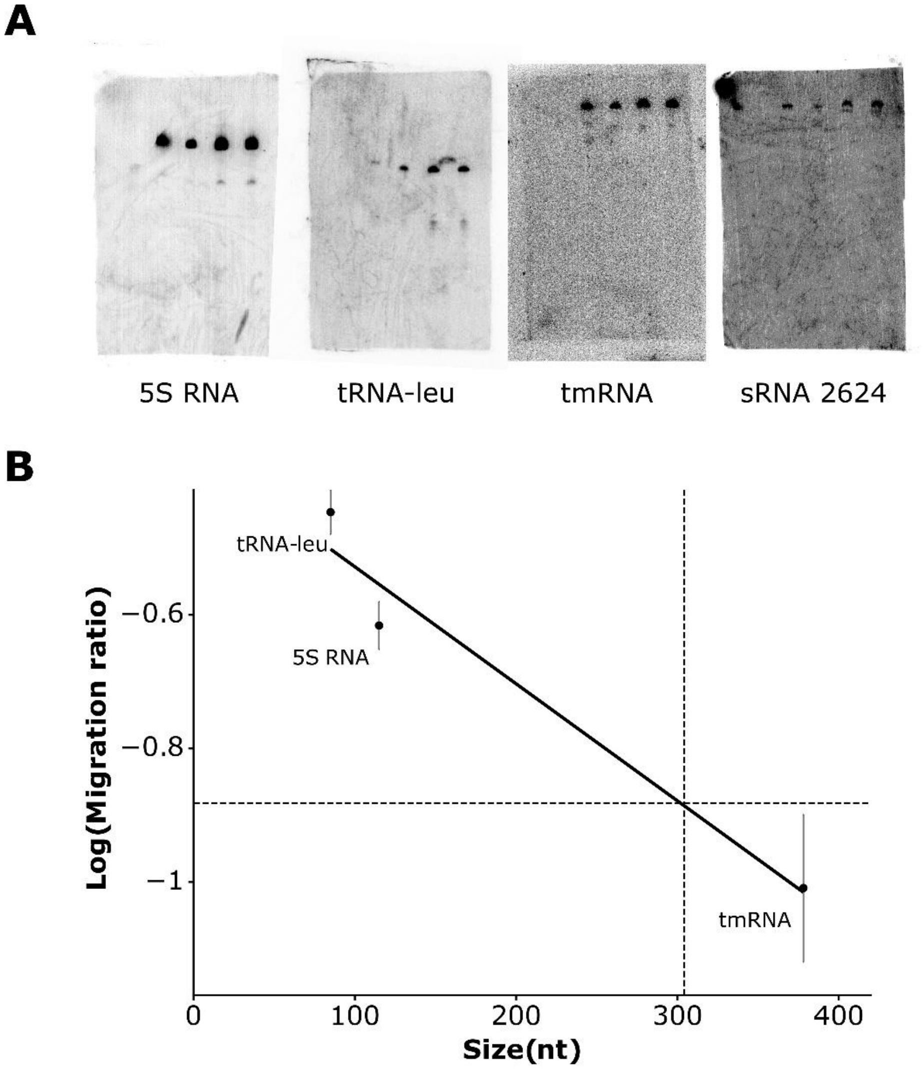

**Figure S1. Size estimation of candidate sRNA2624 based on known RNA**. Hybridization of probes for controlled RNA is observed with radioactivity labelling. Since the size of these controlled RNAs are known (5S RNA, 115 nt; tRNA-leu, 85 nt and tmRNA; 378 nt), their migration ratio in a membrane can be measured and graphed to estimate the size of candidate RNAs such as sRNA2624 (B). All probes were tested on three different membranes.

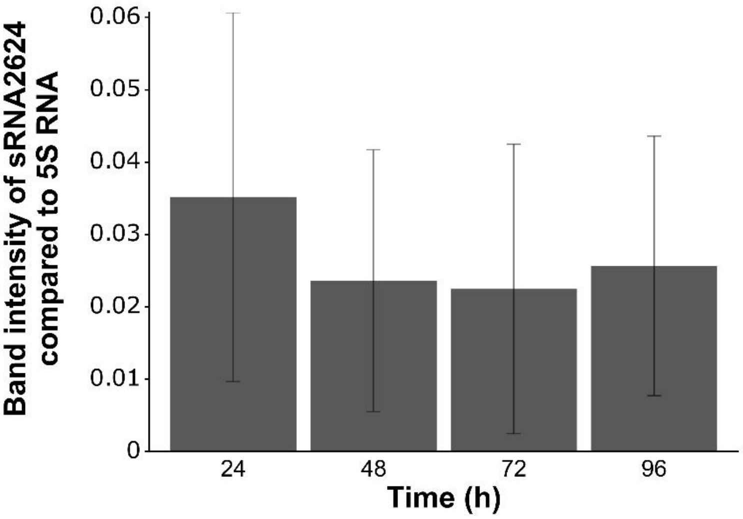

**Figure S2. Expression of sRNA2624 over the growth of *M. extorquens.*** Probes for sRNA 2624 were tested on three different membrane and normalized to the band intensity of a control RNA, 5S RNA.

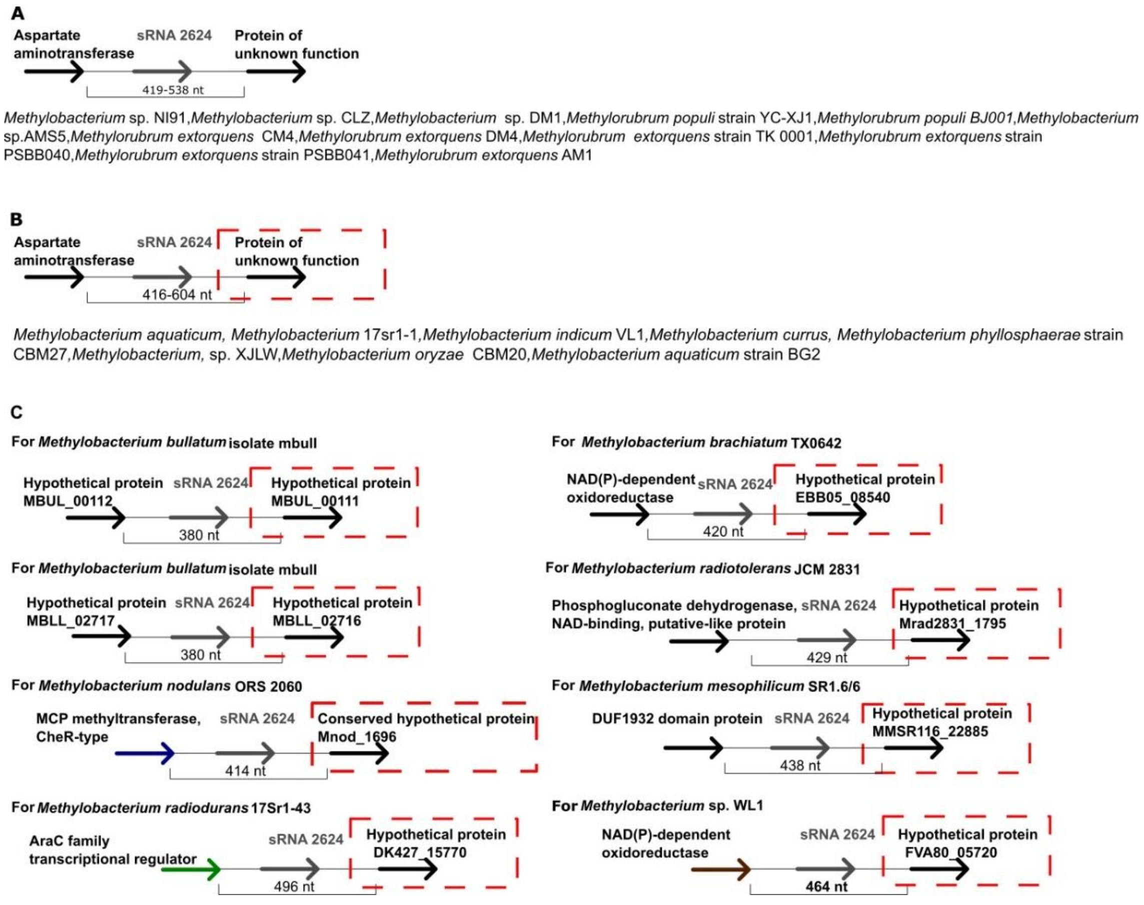

**Figure S3. Genomic context of Met2624.** Results are based on NCBI BLASTn [37]. All bacteria listed in (A) have an aspartate aminotransferase in 5’ of Met2624 and a protein of unknown function of the same family in 3’. Bacteria listed in (B) also have an aspartate aminotransferase upstream of the sRNA candidate, but it is followed by a different protein of unknown function. Bacteria listed in (C) have the same protein of unknown function encoded downstream of Met2624 in their genome as those listed in (B) represented by a dashed red box, but the protein upstream differs. Proteins that are identical are represented by the same color in (C) (black, blue, brown, or green). Met2624 is encoded only in the family Methylobacteriaceae.

```
Sequences: 34
Columns: 200
Reading direction: forward
Mean pairwise identity:  83.71
Shannon entropy: 0.37902
G+C content: 0.72183
Mean single sequence MFE: -111.29
Consensus MFE: -73.21
Energy contribution: -68.23
Covariance contribution:  -4.98
Combinations/Pair:   1.71
Mean z-score:  -2.03
Structure conservation index:   0.66
Background model: dinucleotide
Decision model: sequence based alignment quality
SVM decision value:   1.47
SVM RNA-class probability: 0.941197
Prediction: RNA
```

**Figure S4. RNAz results for intergenic regions containing Met2624.** The program suggested Met2624 is a functional RNA.

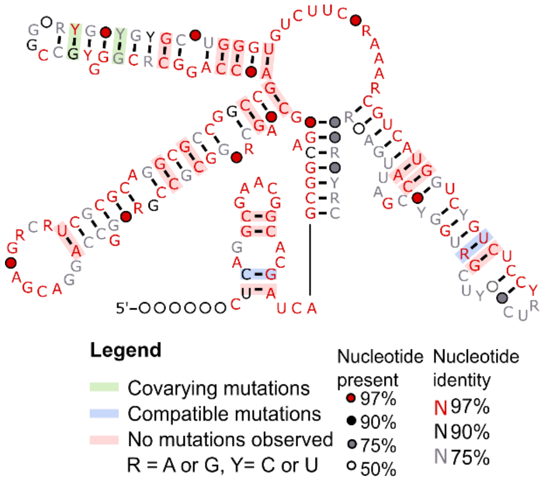

**Figure S5. Secondary structure for Met1153.** The structure was drawn by the program R2R [53]. Taken individually, none of the indicated covarying base pairs are considered statistically significant according to R-scape [44].

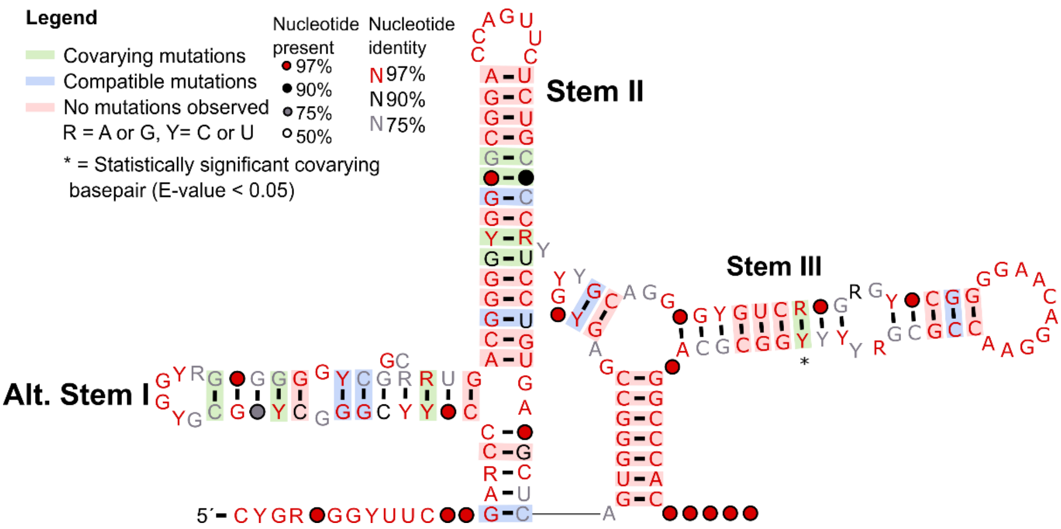

**Figure S6. Alternative secondary structure for Met2624.** The structure was drawn by the program R2R using an alternative Stockholm alignment [53]. Taken individually, a base pair in stem III is considered statistically significant according to R-scape [44]. While the structure pictured in Figure 5 fits the conserved region and the experimentally estimated 5′ and 3′ ends, several alternative alignments (and corresponding structure predictions) with slightly different 5′ and 3′ ends were performed (data not shown). For several of these structures as well as for the structure pictured in Figure S6, it can be noted that stems II and III are identical to that of Figure 5. This contrasts with the “Alternative stem I” which is completely different from Stem I in Figure 5 and also appears less likely to form, given that the latter has more conserved basepairs, covariations and compatible mutations.

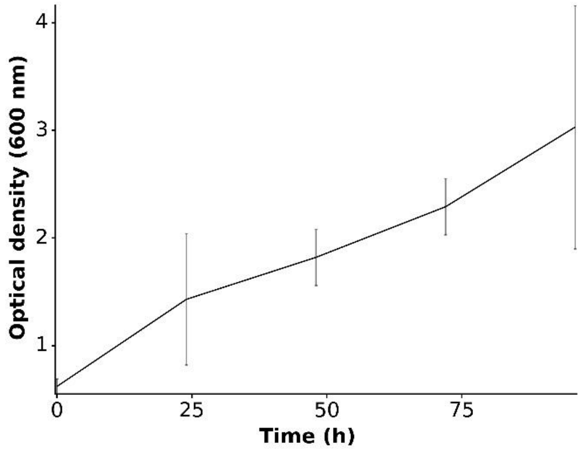

**Figure S7. Growth curve of *M. extorquens* ATCC55366 for Northern Blot Analysis.** It was cultivated with 1% methanol as the sole source of carbon in tri-replicates. The optical density (600 nm) was measured every 24 hours for four consecutive days.
